# When discrete characters are wanting: Continuous character integration under the phylospecies concept informs the revision of the Australian land snail Thersites (Eupulmonata, Camaenidae)

**DOI:** 10.1101/2025.04.17.649343

**Authors:** Guoyi Zhang, Gerasimos Cassis, Frank Köhler

## Abstract

Species that are predominantly characterized by continuous instead of discrete morphological characters pose a challenge to species delimitation. Under the phylospecies concept, species are delimited by apomorphies, which are difficult to establish when characters are not discrete. In the present study, we address these challenges in the Australian land snail genus *Thersites* (family Camaenidae), where accepted species exhibit a scarcity of discrete distinguishing characters and differ in a limited range of continuous characters according to prior taxonomic studies. We integrate analyses of genome-scale molecular data with evaluations of several continuous, qualitative, and discrete morphological characters derived from landmarks to delineate species by identifying apomorphies. We found that dimensions of shell and genitals overlapped considerably among the currently accepted species. These overlaps may indicate morphospace saturation, which may affect species delineation through the gen-morph species concept. Statistical methods, such as Dunn’s test, failed to consistently delineate monophyletic taxa. Additionally, we could not derive apomorphies from discretized landmarks for the prevalence of outlier specimens. However, applying the Kruskal-Wallis test at certain nodes of the tree revealed significant differences in some continuous characters. We propose that these inferred differences represent apomorphies of taxonomic lineages. Ultimately, we suggest based on our findings that of the four currently accepted species, two should be synonymized. To maintain monophyly of taxa, we synonymize *Thersites mitchellae* with *T. novaehollandiae* and *T. darlingtoni* with *T. richmondiana*. Distinct character states of the umbilicus support the existence of two independent lineages: One with an open (*T*. sp1 + *T*. sp2) and one with a closed umbilicus (*T. novaehollandiae* + *T. richmondiana*). Overall, the Kruskal-Wallis tests support four distinct species, *T. novaehollandiae* and *T. richmondiana* plus two newly described species.

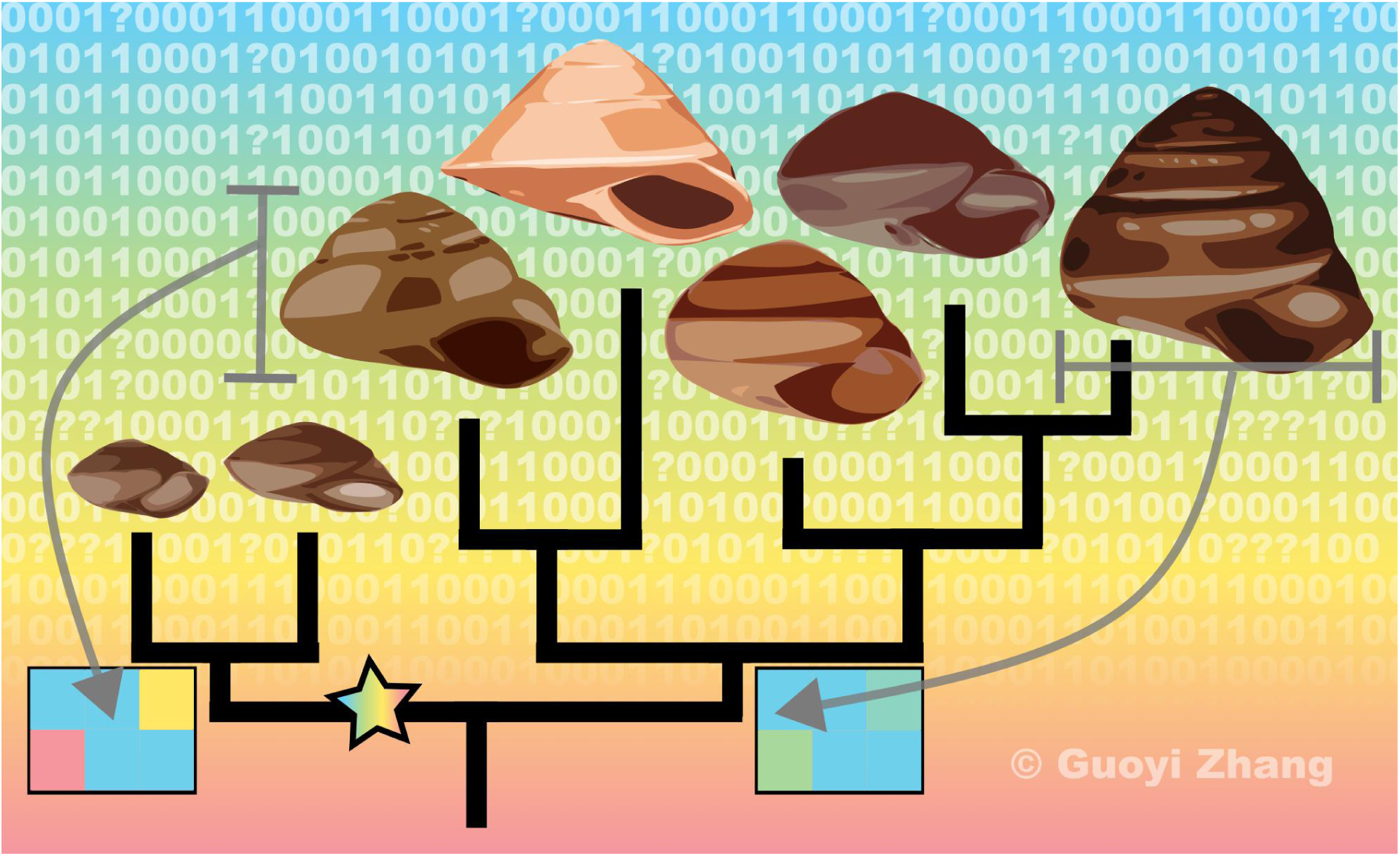

## Introduction

Species delimitation is fundamental to biological and evolutionary studies (Simpson, 1951). Morphological characters are commonly employed for this purpose (Zapata & Jiménez, 2012), and traditionally taxonomists have exclusively relied on morphological traits to define species for a long time indeed (Blackwelder, 1967). While molecular characters have gained broad significance in recent decades, especially in resolving genealogical relationships (Lee and Palci, 2015), morphological characters remain critically important, especially in delimitating taxa. Morphological characters can broadly be categorized as either discrete or continuous. When discrete characters are available, qualitative assessments are typically used in taxonomic delimitation and character optimization (Farris, 1998; Swofford, 2003; Goloboff and Morales, 2023). In contrast, when characters are continuous, statistical methods and phenetic methods are often applied to distinguish character states. However, the statistical approach has proven to be of limited value because character homology is not considered (Sneath and Sokal, 1973).

The challenge of delimiting species is further compounded by the lack of standardized approaches for processing the different types of characters mentioned above. Some authors even proposed to use characters from DNA barcoding to delimit species (Hartop et al., 2022). However, what defines ‘species’ remains a complex debate that cannot be fully resolved by relying on a single character or DNA sequence alone, especially within a cladistic framework (Avise, 2000; Will & Rubinoff, 2004). Moreover, various species concepts differ in how they select, process, estimate, and apply characters. Wilkins (2018) catalogued 28 species concepts, among which the phylospecies concept (Hennig, 1950) is one of the more significant ones (Wilkins, 2018; Odenbaugh, 2022). This concept defined species as the smallest diagnosable monophyletic biological entity identifiable by apomorphies (Hennig, 1950; Wilkins, 2018). However, challenges to the practical application of this concept arise for groups lacking discrete morphological apomorphies even when significant molecular differences may be evident. This issue is exacerbated by the prevalent preference for discrete characters as apomorphies over continuous ones (Rae, 1998). Although efforts have been made to incorporate continuous data into character optimization (Thiele, 1993; Goloboff et al., 2006) and even landmark datasets (Zelditch et al., 1995; Catalano et al., 2010) in systematic studies, such approaches do not seek to establish reliable apomorphies that are derived from continuous characters but merely represent phenetic comparisons.

Integrative species concepts promise to overcome the limitation of artificial species boundaries by combining multiple criteria. For example, Hong (2020) proposed the gen-morph species concept, which builds upon de Queiroz’s (1998) general lineage species concept and integrates several existing approaches to enhance operability. A key requirement of this concept is that a species must exhibit at least two distinct, statistically discontinuous apomorphies. However, when species occupy broad ecological niches with high niche variability while character variation is constrained, a continuous overlap in the morphospace (Budd, 2021) of taxa may occur (Foote, 1994; Paup, 1966; Erwin, 2006). Therefore, the gen-morph species concept is challenged by saturation in morphospace. Moreover, Mayden’s (1997) and Wiley and Lieberman (2011) argued that operational species concepts or species concepts requiring additional criteria ultimately capture only a subset of biodiversity.

The Australian land snail *Thersites* Pfeiffer, 1855 (Camaenidae) exemplifies the challenges of delimiting species under these constraints. The members of this group exhibit predominantly continuous differences in morphological characters while they may also face morphospace saturation due to ecological, functional, and genetic constraints. For example, such morphospace has been attributed to the retention of certain shell morphotypes in phylogenetically distinct camaenids that inhabit xeric environments in northern Australia for selection likely favoured these morphotypes in response to a harsh, yet homogeneous environment (Criscione & Köhler, 2013).

Currently, *Thersites* comprises four accepted species: *Thersites novaehollandiae* (Gray, 1834), *T. richmondiana* (Reeve, 1852), *T. mitchellae* (Cox, 1864), and *T. darlingtoni* Clench & Archer, 1938 (Stanisic et al., 2010). Bishop (1978) reviewed the genus based on comparative morphology using continuous characters derived from the shell and reproductive organs. In this revision, Bishop (1978) treated *T. darlingtoni* as a junior synonym of *T. richmondiana*. Subsequently, Stanisic et al. (2010) treated *T. darlingtoni* as an accepted species for being ‘distinguishable by smaller size and a more rounded body whorl’. As noted above, the lack of discrete diagnostic features hampers the identification of unambiguous apomorphies rendering the current classification phenetic. To complicate matters, a recent molecular phylogenetic study revealed that *Thersites novaehollandiae*, a species characterized by a variable morphology and a wide distribution, is made up of multiple mitochondrial lineages, and is paraphyletic with respect to *T. mitchellae* (Hugall & Stanisic, 2011).

This finding contradicts the earlier taxonomic classification of Bishop (1978) who accepted *T. novaehollandiae* and *T. mitchellae* as two distinct species. The inconsistency between mitochondrial phylogenetics and morphology-based taxon delineation by which two morphologically distinguishable forms are not mutually monophyletic in a phylogenetic tree underscore the broader challenges of delineating species by emphasizing continuous traits.

*Thersites* species inhabit a variety of evergreen to deciduous forest types throughout eastern New South Wales and south-eastern Queensland. However, recent surveys in more xeric landscapes west of the Great Dividing Range have recorded presumedly undescribed species of *Thersites.* These findings imply that this genus may not be limited solely to the mesic forests in central-eastern Australia (Murphy and Shea, 2015, 2023). These potentially new species, *Thersites* sp. “MtKaputar” (Murphy and Shea, 2015) and *Thersites* sp. “Coolah Tops” (Murphy and Shea, 2023), have been discovered in drier sclerophyll woodlands in central-eastern New South Wales. However, the taxonomic status of these two candidate taxa remains to be clarified.

The evolutionary history of *Thersites* is poorly understood, partly because previous studies have not sampled all taxa comprehensively and relied exclusively on mitochondrial DNA. In this study, we employe a genome-wide SNP dataset covering multiple populations across all accepted taxa to test hypotheses concerning the species-level classification as well as the evolutionary history of this group. Given the lack of clearly distinguishing, especially categorical, characters, we aimed to objectively quantify various allegedly diagnostic traits to derive additional categorical characters that could serve as potential apomorphies in species delimitation. These characters include Voronoi distances used to assess soft body features and landmark datasets used to assess shell shapes.

## Materials and Methods

Our methodology pipeline is outlined in the diagram depicted in Figure 1.

**Fig. 1.**
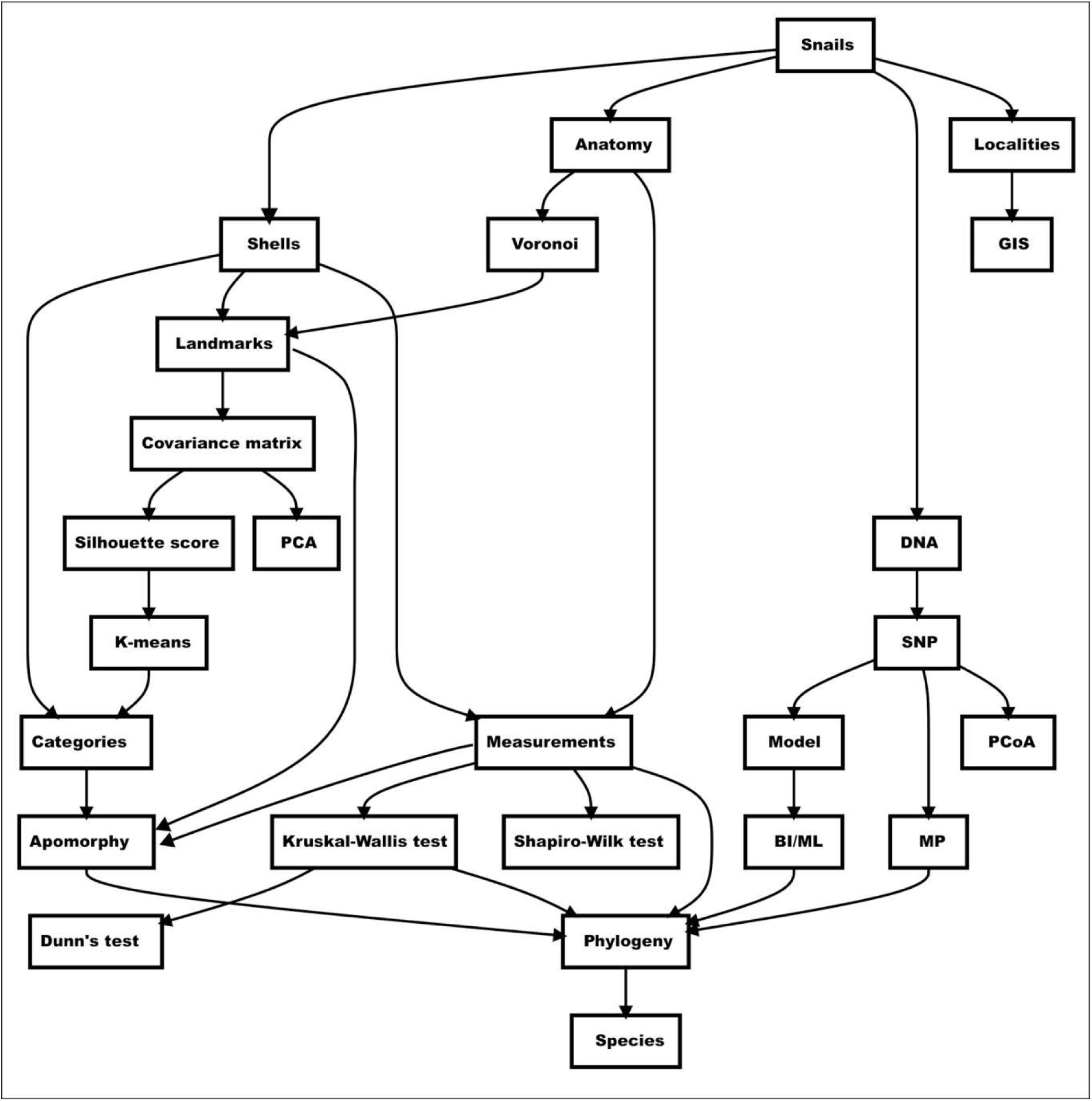
Analytical work flow used in this study.

### Taxonomic classification

The latest taxonomic classification of *Thersites* by Stanisic et al. (2010) serves as our null hypothesis in species delimitation. In addition, our null hypothesis includes the two supposedly undescribed species recognised by Murphy and Shea (2015, 2023).

### Materials examined

Vouchers studied here (for shell morphology, Dryad morphology/shell/measure_shell.csv;; for reproductive anatomy, Dryad morphology/genital/measurement/measure_genital.csv; for sequenced specimens, Dryad character/morphology.csv), including types, are kept in the collections of the Australian Museum, Sydney (AM), Queensland Museum, Brisbane (QM), Museum Victoria, Melbourne (NMV), Natural History Museum, London (NHMUK), and the Museum of Comparative Zoology, Cambridge, Mass. (MCZ).

### Data repository and analytical scripts used

All used scripts used in this manuscript and related instructions have been deposited in the GitHub repository (http://github.com/starsareintherose/Thersites-Data). Datasets are available from Dryad (https://datadryad.org/stash/dataset/ doi:10.5061/dryad.2rbnzs81c). Operations were taken under BioArchLinux environment (Zhang et al., 2025)

### Species occurrences

We obtained available specimens’ GIS information from the Atlas of Living Australia (ala.org.au), which includes records from all Australian museums. We used QGIS to plot all available specimen records on a map, categorized by originally assigned taxon names (Fig. 2). Occurrences of specimens included in the molecular analyses are marked with a star symbol.

**Fig. 2.**
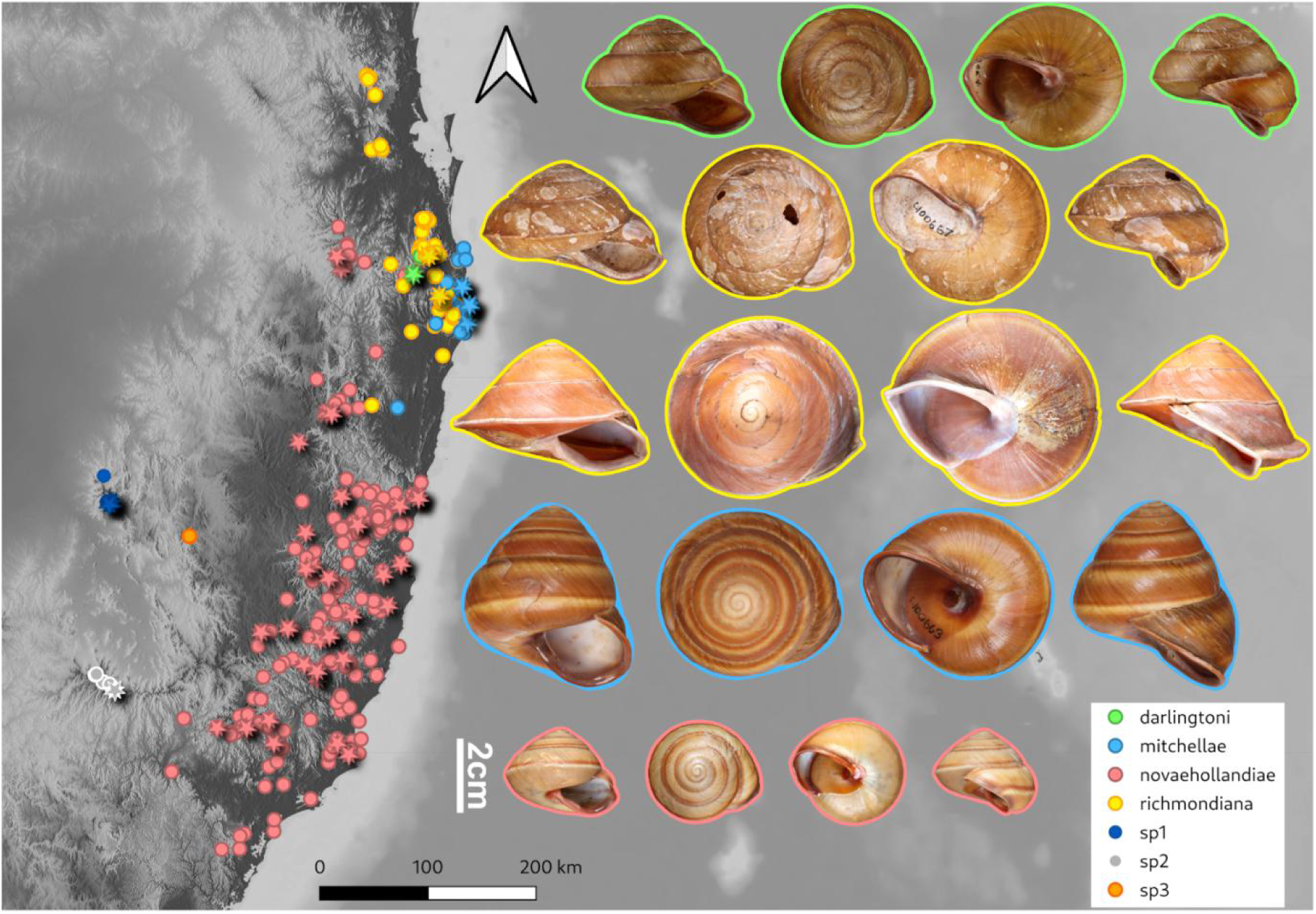
Distribution of *Thersites* species based on the collections of Australian museums. Occurrences of sequenced specimens marked with a star. Type specimens of *Thersites*, from top to bottom: *Thersites darlingtoni*, MCZ 99054; *Annakelea tympanum*, AM C.100667; *Helix richmondiana*, NHMUK 1983079; *Annakelea peragrans*, AM C.100663; *Helix dupuyana*, NHMUK 1977032.

### Shell measurements

Shell dimensions of 323 adult specimens were measured and analysed to test if the shell width–height ratio is a suitable characteristic to distinguish species. We applied focus stacking to generate high-resolution images. Subsequently, the shells were extracted from the background and resized to a uniform pixel scale using Adobe Photoshop 2024 or GIMP 2.10 (GIMP Development Team, 2025) with G’MIC-Qt plugin (Tschumperlé et al., 2025) Next, we converted all images to a white-shell and black-background format using ImageMagick (ImageMagick Studio LLC, 2025). Finally, we utilized a custom FIJI (Schindelin et al., 2012) macro script to measure shell width and height, which were then visualized as scatter plots using matplotlib (Hunter, 2007) implemented in Python (Python Software Foundation, 2025).

After excluding broken shells unsuitable for morphometric analysis, we conducted a landmark-based morphometric analysis on 321 aperture view photographs to assess shell shape variation. This decision followed Stone (1998)’s opinion considering the homology of landmark assignments. Stone (1998) showed that the aperture view is most useful to describe and analyze shell shapes. Seven fixed landmarks: LM1, shell apex (embryonic shell); LM2, the left intersection point of the penultimate whorl and its preceding whorl; LM3, the left intersection point of the body whorl and penultimate whorl; LM4, left edge point of callus and aperture; LM5, right intersection point of aperture and body whorl; LM6, the right intersection point of the body whorl and penultimate whorl; LM7, the right intersection point of the penultimate whorl and its preceding whorl.These seven landmarks and two semi-landmark curves were digitized using tpsDig 2.32 (Rohlf, 2021a). Fifty equidistant semi-landmarks were samples along each curve. Along with the landmarks, these semi-landmarks were treated equally in the analyses. One curve traces the left outline of the body whorl and the other delineates the aperture shape and the seven landmarks, whose positions are shown in Fig. 3A. We then analyzed three sets of landmarks or semi-landmarks separately: the two curves as distinct groups and the seven landmarks as a single group. For each individual group and the combined dataset, we performed morphometric analyses independently. In each analysis, we applied a Procrustes fit by aligning the principal axes to mitigate potential manual rotation errors. Covariant distance matrices and principal component analysis (PCA) were then computed using MorphoJ 1.08.02 (Klingenberg, 2011), and PCA scatter plots with convex hulls were visualized. We calculated the average observations for the initial hypothesis and visualized them using the thin plate spline network method in MorphoJ. We then performed a discriminant function analysis in MorphoJ to test the difference between two groups of specimens using 1,000 permutation runs. Average correction rates of cross-validation were calculated by a custom python script. Following these separate analyses, we examined the scatter plots to ensure that the negative control specimens were not completely intermixed with the other groups. Here we considered the keeled shell, *Thersites richmondiana*, as a negative control. If all other groups largely overlapped with this negative control group, the tested landmark group was removed. After removing the landmark group, we combined all points and repeated the analysis.

**Fig. 3.**
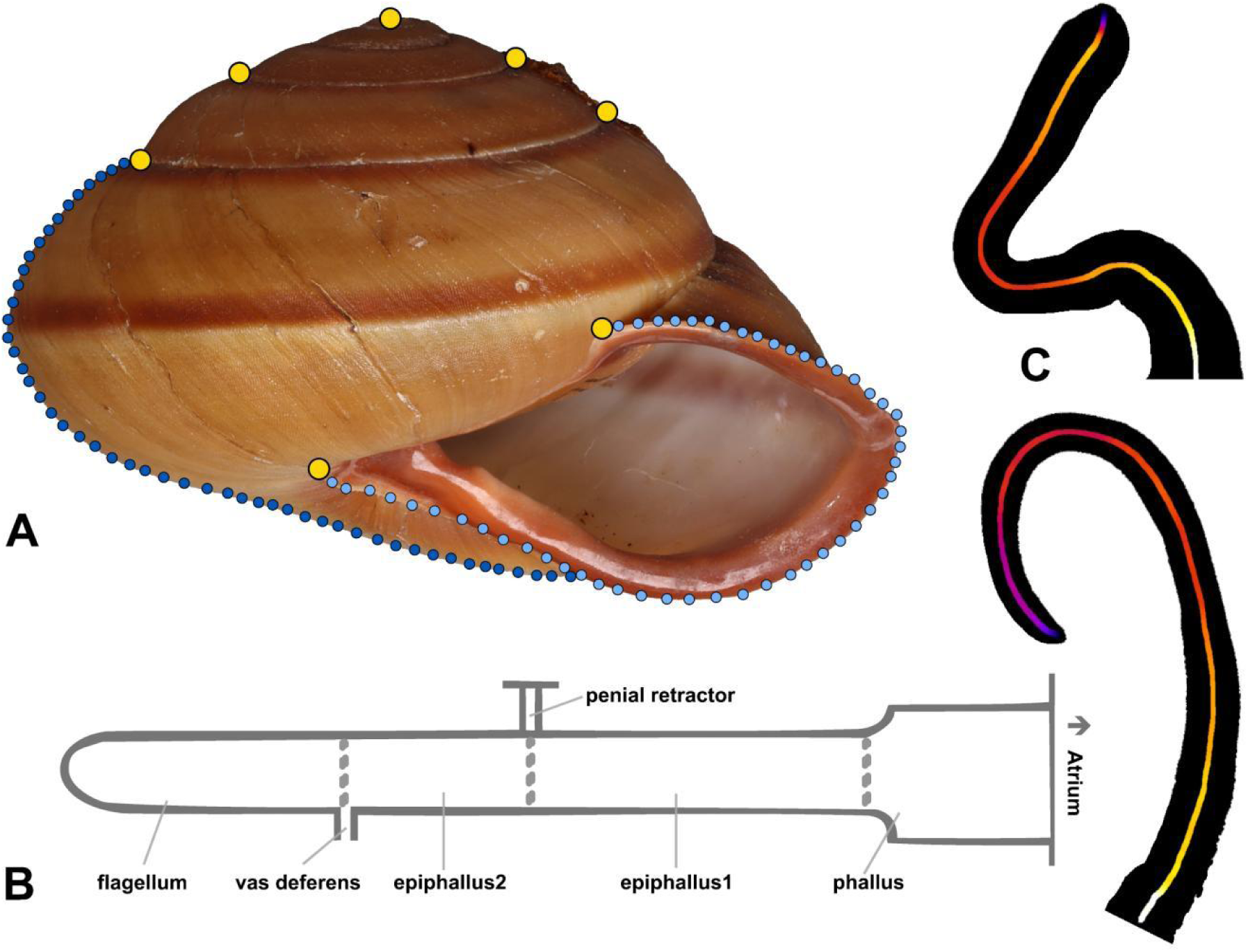
A. Landmarks used for morphometric geometric analyses. Yellow dots: Landmarks; dark blue dots: Semi-landmark curve 1 along periphery of body whorl; blue dots: Semi-landmark curve 2 along apertural lip. B. Diagram of genital terminology. C. Diagram of the flagellum width changes along the middle line based on Voronoi distance. The heatmap follows white (thick) - yellow – red – purple (thin).

### Genital system measurements

We measured the genital systems of 89 specimens using FIJI (Schindelin et al., 2012), separately quantifying the lengths of phallus, epiphallus, and flagellum. Epiphallus1 and epiphallus2 are delimited by the junction with the penial retractor muscle (Fig. 3B). Epiphallus1 is the part of the epiphallus proximal and epiphallus2 distal of that junction (Fig. 3B). Measurements of genital organs are visualized using histograms. We performed Shapiro-Wilk’s test to assess the normality of the values obtained. The proximal epiphallus to the flagellum ratio was then visualized in violin maps and scatter plots. We tested Bishop’s (1978) hypothesis that species differ in the relative lengths of their genital organs by analysing the ratios between the lengths of the proximal ephiphallus, whole phallus, epiphallus1, phallus + epiphallus1, phallus + epiphallus, respectively. Additionally, we plotted the epihallus1 – epiphallus ratio in a violin map and scatter plots. We performed a Kruskal–Walli’s test to assess significant differences among the groups, followed by Dunn’s test as a post-hoc analysis to determine the specific groupings based on the Kruskal–Wallis results.

We evaluated the flagellum shape using width measurements and landmark-based morphometrics (Fig. 3C). After discarding incomplete or broken flagellum unsuitable for analysis, candidate photographs were standardized to a uniform pixel scale and converted to a white flagellum on a black background using ImageMagick. Voronoi distances were computed using FIJI and the width profile tools, and the resulting flagellum shape metrics for 81 specimens were analyzed. After collecting the Voronoi distance data, we used a custom Python script to resample the distance points to 100 points per specimen. We analysed three different datasets corresponding to each 100%, 75% and 50% of flagellum length measured from the tip to interrogate the relevance of flagellum tip shape over the shape across the entire length of the flagellum. We visualized the flagellum shape curves using matplotlib in Python and converted the resampled points to TPS files via a custom Python script. Subsequently, all landmarks were analyzed in MorphoJ, where we performed a Procrustes fit, computed covariant distance matrices, and conducted a PCA. We also calculated the average observations by group. Finally, PCA results were visualized using scatter plots and convex hulls, and group average observations were displayed using the thin plate spline network. Additionally, we also performed a discriminant function analysis in MorphoJ to test the difference between two groups of specimens using 1,000 permutation runs. Average correction rates of cross-validation were calculated by a custom python script.

### SNP generation and analyses

DNA extracts from 80 individuals were used to perform DArTseq high-density genotyping (DArT Pty Ltd., Canberra, Australia), a Single-nucleotide polymorphism (SNP) based population genetic method (Melville et al., 2017). The workflow of this commercial application includes a double restriction enzyme-mediated genome complexity reduction method to target multiple loci that are randomly distributed across the genome and uses next-generation sequencing platforms to sequence the resulting DNA fragments (Melville et al., 2017). For our current dataset, the PstI/HpaII enzyme combination was employed to generate short DNA fragments based on preliminary trials testing various restriction enzyme combinations to optimize genome representation, which were sequenced as single-ended reads on an Illumina Hiseq 2500 platform. These reads were processed using a proprietary DArT analytical pipeline (Jaccoud et al. 2001) to obtain a SNP matrix for subsequent analyses.

We used the dartR package (Mijangos et al., 2022) for the R statistical environment (R Core Team, 2025) to process and analyse the SNP matrix. Initially, we filtered the data as follows: exclude loci with a read depth outside the 5–30, removal of markers with reproducibility scores below 100%, exclude loci with coverage of below 80% of all individuals, and excluding individuals with coverage of below 50% of all loci, removal of all monomorphic markers. We also carried out quality control by filtering out SNPs with minor allele frequencies below 0.01 and randomly removing secondary markers to mitigate potential linkage following the dartR tutorials. We calculated the Principal Coordinates Analysis (PCoA) to visualize the genetic structure using ggplot2 (Wickham, 2009).

### Molecular phylogenetic analyses

We analysed the SNP data matrix to reconstruct the phylogenetic relationships among the sequenced specimens. To thi send, we reconstructed Maximum Parsimony (MP) phylogenies using TNT 1.6 (Goloboff and Morales, 2023) with guoyi.run (Zhang, 2025) under three weighting schemes: Equal weighting, implied weighting, and extended implied weighting. For implied weighting and extended implied weighting, we set the inital K value to 12 following the recommended K for morphological character matrix from Goloboff et al. (2017). The guoyi.run script performed 1,000 iterations of TBR branch swapping to generate Maximum Parsimony Trees (MPTs), which were subsequently summarized using a strict consensus tree. Group support was estimated using 1,000 jackknifing iterations and 1,000 symmetric resampling replicates. We used the default deletion probability of 36 for jackknifing and symmetric resampling. Additionally, we calculated the consistency index (CI), retention index (RI), and tree length (TL) using guoyi.run. After initially performing EIW and IW analyses with K = 12, we further explored K values ranging from 1 to 100, performing 10 cycles of ratchet and drift followed by branch swapping for each K. For EIW and IW, we then selected the best K based on CI, RI, and TL. In cases where multiple K values performed equally well, we summarized the resulting trees using a strict consensus approach.

For model-based phylogenetic analyses, we employed modeltest-ng (Darriba et al., 2020) to identify the most suitable substitution model based on the Bayesian Information Criterion (BIC), incorporating 16 gamma categories. The appropriate model was implemented in a Maximum Likelihood (ML) analysis performed by using raxml-ng (Kozlov et al., 2019). We utilized 10 parsimony starting trees and 10 random starting trees and applied the SPR branch swapping to obtain the best Maximum Likelihood tree. Group support was estimated by performing 1,000 Felsenstein bootstrap replicates by raxml-ng and 1000 jackknifing replicates with 50% subsampling proportion by IQ-Tree 3 (Wong et al., 2025). We reconstructed a Bayesian Inference (BI) phylogeny by using MrBayes 3.2.7 (Ronquist et al., 2012) by applying default prior, running two independent runs for 2,000,000 generations, sampling every 2,000 generations, with a chain temperature of 0.05. The final consensus tree and posterior probabilities were determined from the last 50% of all sampled trees (burn-in = 50%).

ML bootstrap (BS), MP jackknifing (PJK) and likelihood jackknifing (LJK), symmetric resampling (SR) values above 90% and Bayesian posterior probabilities (BPP) above 0.9 are considered to indicate strong support; values of BS/JK/SR >70% or BPP >0.8 indicate good support; lower values are regarded as insufficient support.

Phylogenetic networks were inferred using a parsimony-based approach implemented via the InferNetwork_MP command in PhyloNet 3.8.4 (Yu et al., 2013). The input comprised phylogenetic tree inferred from Bayesian methods. To identify the optimal network, a steepest descent search algorithm was applied. For visualization, the five inferred networks were summarized in SplitsTree6 (Huson & Bryant, 2024) using the consensus outline method (Huson & Cetinkaya, 2023), yielding a representative network.

### Morphological character optimization analysis

We used Wincladtree (Goloboff, 2024) with TNT to visualize the continuous apomorphic character along the molecular phylogenetic tree. The continuous characters include shell width and height, lengths of phallus, epiphallus1, epiphallus2, and flagellum.

We used phytools (Revell, 2024) to visualize continuous measurements along the phylogenetic trees. We mapped continuous character states relating to shell width, shell height, lengths of phallus, epiphallus1, epiphallus2, and flagellum onto the molecular phylogeny using the ML estimates under a Brownian evolutionary process. We estimated missing data using ML estimations.

We applied a Kruskal-Wallis test to determine whether continuous dimensions significantly differed among taxa, and the results were visualized using heatmaps. Our analysis aggregated data from descendant taxa at internal nodes and calculated p-values for each variable. If any variable has an empty group (i.e., no valid data), the test is skipped, and the p-value is assigned as n/a (not applicable).

We used the nts matrix files generated from tpsRelw (Rohlf, 2021b) alongside raw landmark matrix data to categorize the landmarks as discrete characters. For the raw landmarks (tps dataset), we additionally applied Generalized Procrustes Analysis (GPA) before generating a matrix to maintain consistency with the analyses conducted in MorphoJ. Silhouette scores were first computed for the variation matrices obtained from landmarks and nts to determine the optimal number of clusters (best K value). We then applied K-means clustering with this optimal K to assign specimens into K categories. This categorization was performed separately for four landmark groups: shell landmarks, body whorl semi-landmarks, aperture semi-landmarks, and half-flagellum landmarks.

Finally, after categorizing both the landmark and a open or closed umbilicus, we mapped these characters onto the phylogenetic trees to identify apomorphic and homoplastic character states. This was achieved using three character-optimization methods, fast optimization (ACCTRAN), unambiguous optimization, and slow optimization (DELTRAN), as implemented in WinClada (Nixon, 2021).

## Results

### Species distributions

*Thersites* species are primarily found in the forests of eastern New South Wales (NSW) and southern Queensland in eastern Australia (Fig. 2). *Thersites novaehollandiae* is the most widely distributed species, ranging from Newcastle north of Sydney to Coffs Harbour in eastern NSW. Its range extends inland to include the Gibraltar and Main Ranges. *Thersites mitchellae* occurs in coastal areas from Lismore in northern NSW to the Sunshine Coast in southern Queensland, while *T. richmondiana* inhabits rainforests further inland between Lismore and Gold Coast in Queensland. *Thersites darlingtoni* is restricted to parts of the Border Ranges (NSW-Queensland border), *T*. sp1 is confined to Mount Kaputar, NSW, and *T*. sp2 to the Coolah Tops, NSW. *T*. sp3 is confined to the Warrabah National Park, NSW. The sequenced samples cover the known distributions of all species well except *T. richmondiana* (Fig. 2). Here, we didn’t have any sequence or samples of *T*. sp3.

### Shell dimensions

Shell widths of all measured specimens ranged from 2.2 to 8.2 cm, with an average of 4.9 ± 0.9 cm. Shell heights varied from 1.6 to 6.4 cm, averaging 4.0 ± 0.7 cm. The width-to-height ratio ranged from 0.83 to 1.18, with an average of 0.83 ± 0.12. *Thersites* sp1 and sp2 were notably smaller than all other species (Fig. 4; Fig. S1; Dryad, morphology/shell/grouped_data.csv). The two thresholds in height-to-width ratio of 0.7 and 0.8 given in taxonomic literature did not delineate the morphospace of any taxon (Fig. S2).

**Fig. 4.**
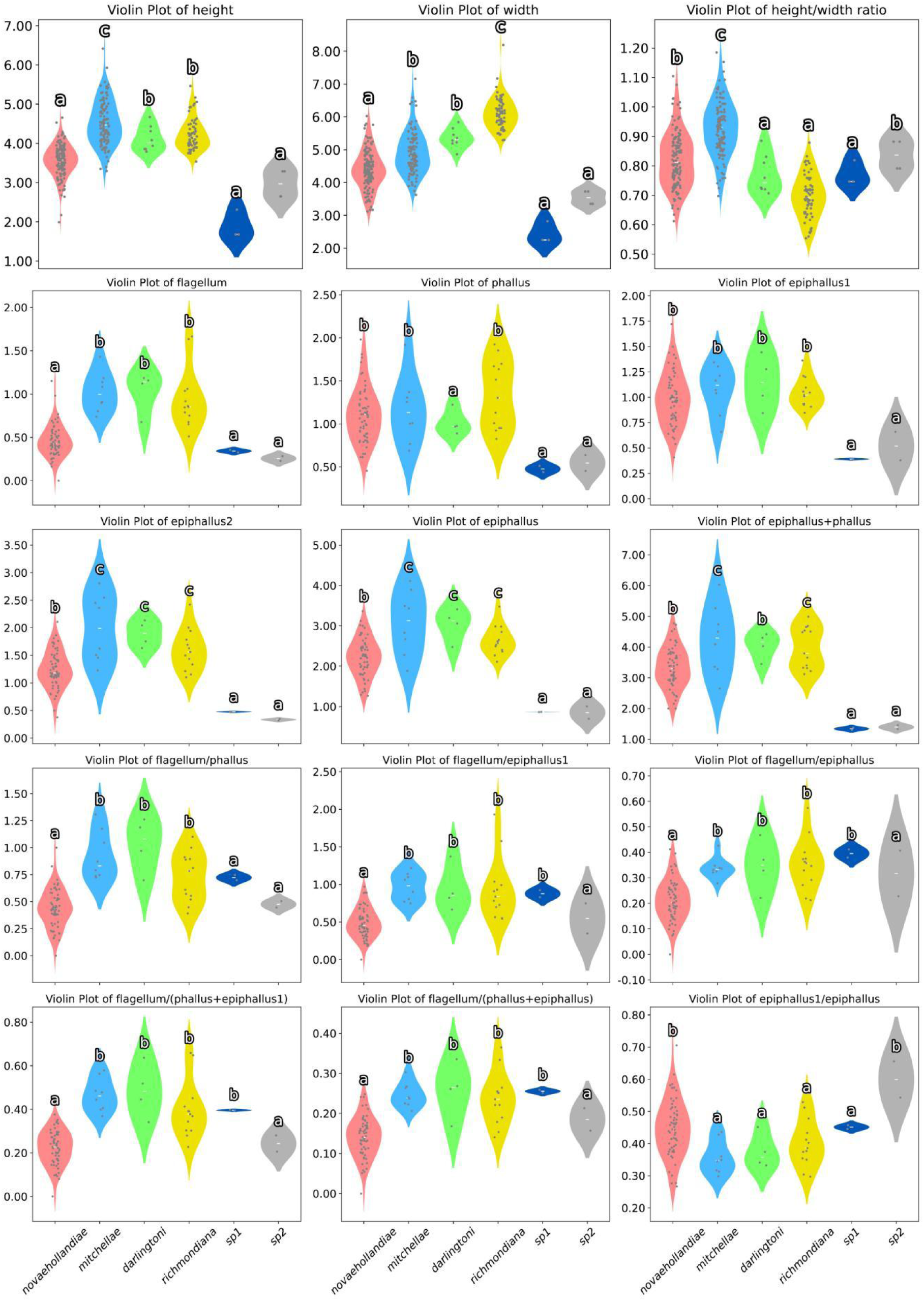
Violin maps of shell height, width, height/width ratio and genital dimensions and ratios. Dunn’s test results are indicated by letters on top of violin maps.

The Shapiro-Wilk test suggested that shell height (W=0.985, p=0.002) and width measurements (W=0.989, p=0.013) did not follow normal distributions; however, the height-to-width ratio did (W=0.993, p=0.176>0.050) (Fig. S1, Dryad, morphology/shell/normality_test_results.txt). Dunn’s test (Fig. 4) indicated that *T. novaehollandiae*, *T*. sp1, and *T*. sp2 consistently clustered together based on these dimensions. When grouped by shell height, *T. darlingtoni* and *T. richmondiana* formed one group while *T. mitchellae* formed a separate group. When grouped by shell width, *T. mitchellae* and *T. darlingtoni* clustered together while *T. richmondiana* occupied a distinct range. Finally, based on the shell height-width ratio, *T*. sp1, *T. darlingtoni*, and *T. richmondiana* grouped together, *T.* sp2 and *T. novaehollandiae* formed a second group, and *T. mitchellae* the third.

### Shell landmark-based morphometrics analyses

In the seven landmarks PCA (Fig. 5A), PC1 accounted for 55% of the variation and PC2 for 20%. The scatter plots showed that *T. richmondiana* overlapped largely with *T. mitchellae* (average accuracy rate [AAR] = 52%) and *T. darlingtoni* (AAR = 93%), and partially with *T. novaehollandiae* (AAR = 53%). Both, *T*. sp1 and *T*. sp2, fell within the range of *T. novaehollandiae* (AAR = 96% and 98.5%).

**Fig. 5.**
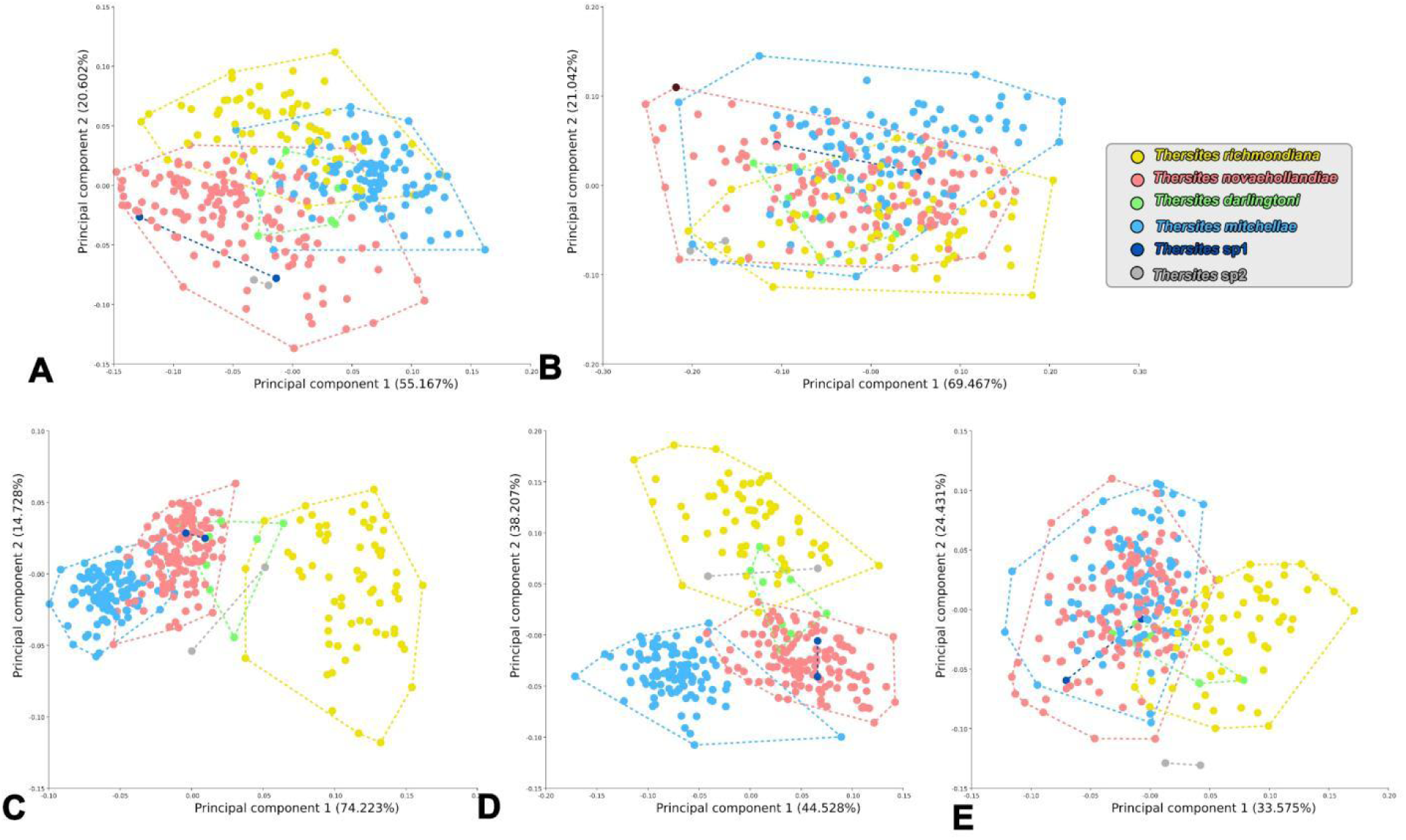
Scatter plots of principal component analysis for seven landmarks (A), semi-landmark curve 2, aperture shape, (B), semi-landmark curve 1, body whorl shape (C), semi-landmark 1 and landmarks (D), two semi-landmark groups and landmarks (E).

In the PCA of body whorl curve landmarks (Fig. 5C), PC1 explained 74% of the variation and PC2 accounted for 15%. The scatter plots indicated that *T. richmondiana* can clearly be distinguished from any other species, except for some overlap with *T. darlingtoni* (AAR = 79.5%) and *T.* sp2 (79.5% AAR with *T. darlingtoni*, 94% with *T.* sp1, others’ AAR are over 98.5%). *Thersites mitchellae* and *T. novaehollandiae* largely overlapped (AAR = 93%), while *T. darlingtoni* overlapped with both *T. novaehollandiae* (AAR = 92%) to some extent as well as in part with *T. richmondiana*. Additionally, *T.* sp1 fell entirely within the range of *T. novaehollandiae* (AAR 95.65%), whereas *T*. sp2 was similar to *T. darlingtoni* (AAR 80%) and *T. richmondiana* (AAR 94%).

For the aperture curve landmarks PCA (Fig. 5B), PC1 explained 69.5% of the variation and PC2 21%. The scatter plots revealed that no species could be distinguished from any other species, as all species overlapped substantially with one another (only the AAR between *T.* sp2 and *T. mitchellae / novaehollandiae* are over 97%, other AAR are lower than 95%).

In the seven landmarks and two semi-landmark curves PCA (Fig. 5E), PC1 explained 23.6% of the variation and PC2 explained 24.4%. Here, *T. richmondiana* overlapped almost entirely with *T. darlingtoni* (AAR 87.7%) and in parts with *T. novaehollandiae* (AAR 98%) and *T. mitchellae* (AAR 99.4%). *Theristes mitchellae* and *T. novaehollandiae* overlapped almost completely with each other (95.9%), while *T.* sp1 fell within their range (with *T. mitchellae*, 98.2%; with *T. novaehollandiae*, 91.3%). Notably, *T.* sp2 could be distinguished from the other species (with *T. novaehollandiae*, 98.5%; with *T.* sp1, 75%, with *T. darlingtoni,* 90%; others over 98.5%).

After removing the aperture curve landmarks PCA (Fig. 5D), which showed negative control mixing with other groups, the PCA yielded PC1 explaining 44.5% of the variation and PC2 explained 38.2%. The scatter plots indicated that *T. richmondiana* partially overlapped with *T. darlingtoni* (AAR 86.3%). *Thersites mitchellae* and *T. novaehollandiae* showed small overlap (AAR 96.7%), whereas *T. darlingtoni* exhibited partial overlap with *T. novaehollandiae* (AAR 88.2%). *Thersites* sp1 fell within the range of *T. novaehollandiae* (AAR 97.1%), and *T.* sp2 within the range of *T. richmondiana* (AAR 97%).

Thin plate spline network maps (Figs 6) revealed that *T. richmondiana* and *T. darlingtoni* tended to exhibit a sharper keel on the body whorl, a more depressed body whorl, a narrower aperture opening, and an overall higher shell spire. By contrast, the maps for *T. mitchellae* and *T. novaehollandiae* indicated a rounder body whorl and a larger aperture. However, *T. mitchellae* tended to have a more elevated spire, while *T. novaehollandiae* typically had a lower spire. Furthermore, *T.* sp1 showed a trend towards a larger aperture, a rounder body whorl, and a lower spire, whereas *T.* sp2 tended to have a narrower aperture, a sharper body whorl, and a lower spire.

**Fig. 6.**
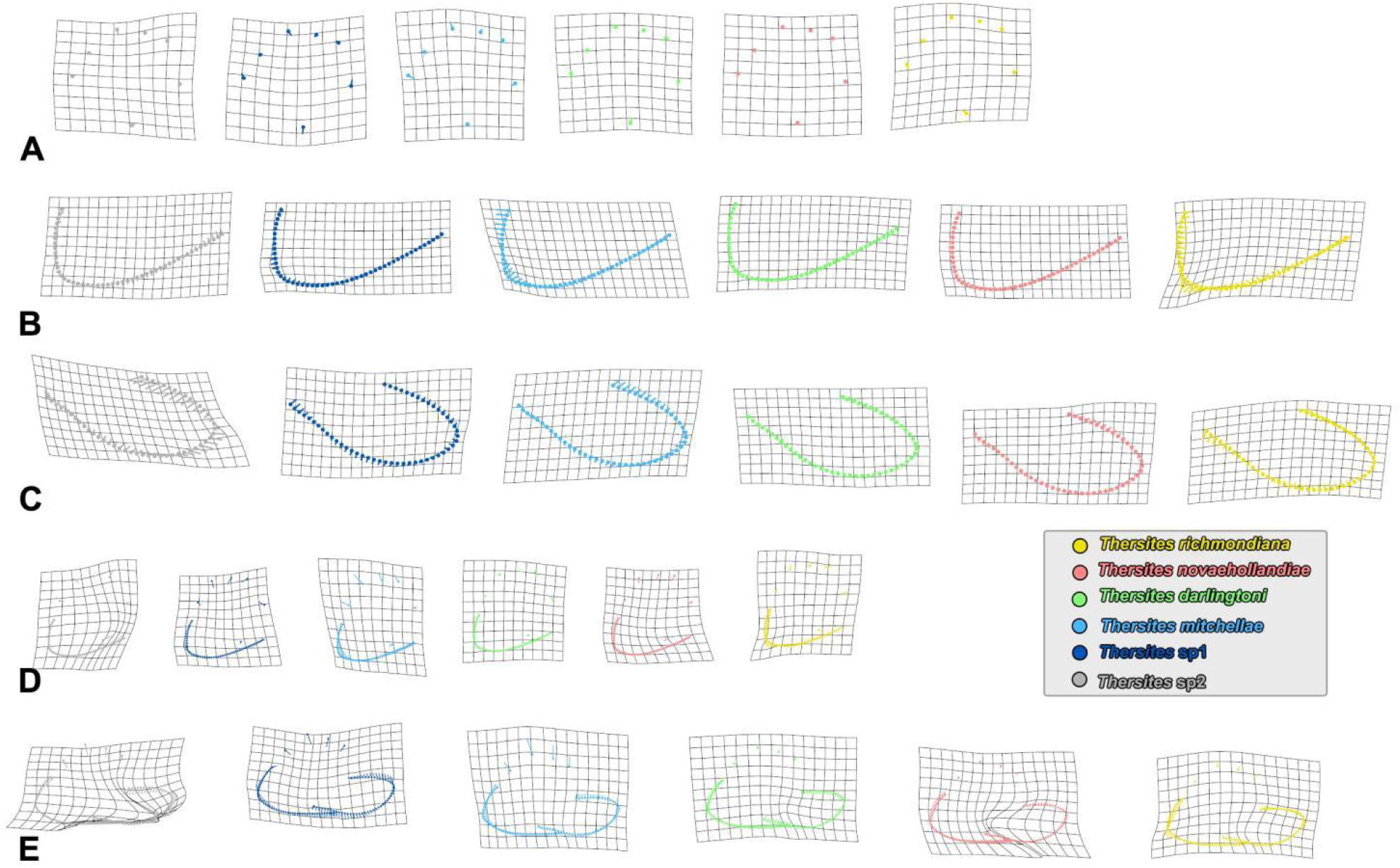
Thin plate spline grid map for each species. A. Seven landmarks. B. semi-landmark curve 1, body whorl shape. C. semi-landmark curve 2, aperture shape. D. semi-landmark 1 and landmarks. E. two semi-landmark groups and landmarks.

### Genital system measurements

Histograms (Fig. S1) and the Shapiro-Wilk test both indicated that the lengths of various genital organs were normally distributed (phallus, W=0.974, p=0.072; epiphallus1, W=0.991, p=0.820; epiphallus2, W=0.976, p=0.095; epiphallus, W=0.988, p=0.614; epiphallus+phallus, W=0.986, p=0.461), except for the flagellum (W=0.913<0.950, p=0.000<0.050). Violin plots combined with Dunn’s test results (Fig. 11) revealed significant differences in genital system dimensions among the groups. Based on flagellum length, *T. mitchellae*, *T. darlingtoni*, and *T. richmondiana* formed one group, while *T. novaehollandiae*, *T.* sp1, and *T.* sp2 formed another. Phallus measurements differentiated into two distinct clusters, one consisting of *T.* sp1, *T.* sp2, and *T. darlingtoni*, and the other of all remaining species. Similarly, the length of epiphallus1 clearly separated *T.* sp1 and *T.* sp2 from all other species. Moreover, both the lengths of ephallus2 and ephiphallus distinguished three groups: *T.* sp1 and *T*. sp2, *T. novaehollandiae*, and all remaining species. The analyses of the epiphallus and phallus data suggested that *T. novaehollandiae* grouped with *T. darlingtoni*, *T. mitchellae* clustered with *T. richmondiana*, and *T*. sp1 and *T*. sp2 consistently formed a separate group.

In contrast, ratio-based analyses (Fig. S) were less discriminative than absolute measurements. The ratios of flagellum/epiphallus displayed a pattern, grouping *T. novaehollandiae*, *T*. sp1, and *T*. sp2 together. Likewise, the ratios of flagellum/epiphallus1, and flagellum/(phallus+epiphallus1), flagellum/epiphallus, flagellum/(phallus+epiphallus), and epiphallus1/epiphallus indicated that *T.* sp2 and *T. novaehollandiae* clustered together, with the other species forming a separate group.

### Flagellum shape variation analysis

Shape analyses of the flagellum based on 100%, 70% and 50% of flagellum length measured from the tip produced inconsistent results. Species could not be distinguished based on analyses of 100% and 70% of the flagellum length. However, analyses based on the distal half of the flagellum (‘50%’) revealed that the tips of the flagellum in *Thersites mitchellae*, *T. novaehollandiae*, *Thersites* sp1, and *T.* sp2 were slimmer than those in *T. darlingtoni* and *T. richmondiana*. The latter two species had a thick flagellum with a tapering tip.

In the Principal component analysis (PCA) based on 100% flagellum length, PC1 accounted for 38.5% of the variation and PC2 explained 24.7% (Fig. S4). *Thersites novaehollandiae* encompassed nearly all the variation in *Thersites* flagellum shape, clustering tightly together (with T. sp1, 96.4%; with others, lower than 90%). Similar patterns have been observed in PCA based on 70% flagellum length (PC1: 39.5%, PC2: 30.2%; Fig. S5) and 50% flagellum length (PC1: 62.2%, PC2: 10.8%; Fig. 7). For 70% flagellum length, the average correction rates were lower than 90%. For 50% flagellum length, AAR were lower than 97%. Scatter plots for 50% flagellum showed that most *T. novaehollandiae* individuals clustered with *T. michellae* (AAR 72.6%), *T.* sp1 (AAR 92.7%), and *T.* sp2 (AAR 96.4%), including one individual of *T. darlingtoni* (AAR 70.7%). Meanwhile, *T. richmondiana* and the majority of *T. darlingtoni* individuals grouped together (AAR 62.5%), even incorporating some *T. novaehollandiae* specimens (AAR 70.7%).

**Fig. 7.**
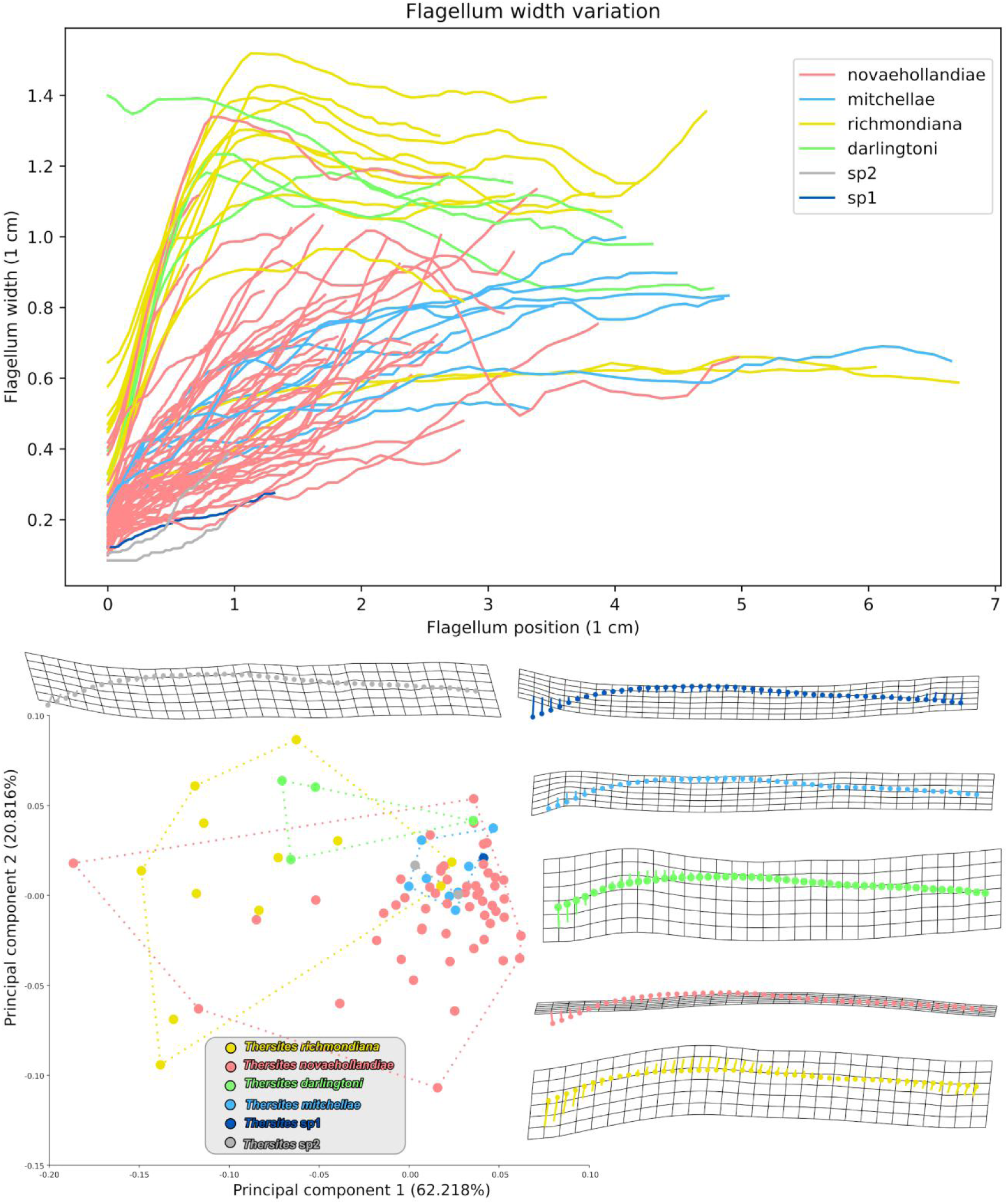
Shape curves and scatter plots of principal component analysis for 50% flagellum length with thin plate spline grid maps for each species.

### SNP analysis

The initial SNP dataset included 80 genotypes with a total length of 103,926 SNPs. After filtering, 73 genotypes containing 765 SNPs remained. The underlying multiple sequence alignment contained 52,408 bp, including on average 10% of missing data. In the PCoA scatter plot (Fig. 8), PC1 accounted for 33.3% of the variation and PC2 for 12.2%. *Thersites novaehollandiae*, *T. mitchellae*, *T. darlingtoni*, and *T. richmondiana* clustered closely along PC1, whereas *T.* sp1 and *T.* sp2 were more distinctly separated on this axis. Notably, some individuals of *T. darlingtoni* grouped with *T. richmondiana*. *Thersites mitchellae* and *T. novaehollandiae* were positioned near each other with *T. mitchellae* exhibiting considerable variation along PC2.

**Fig. 8.**
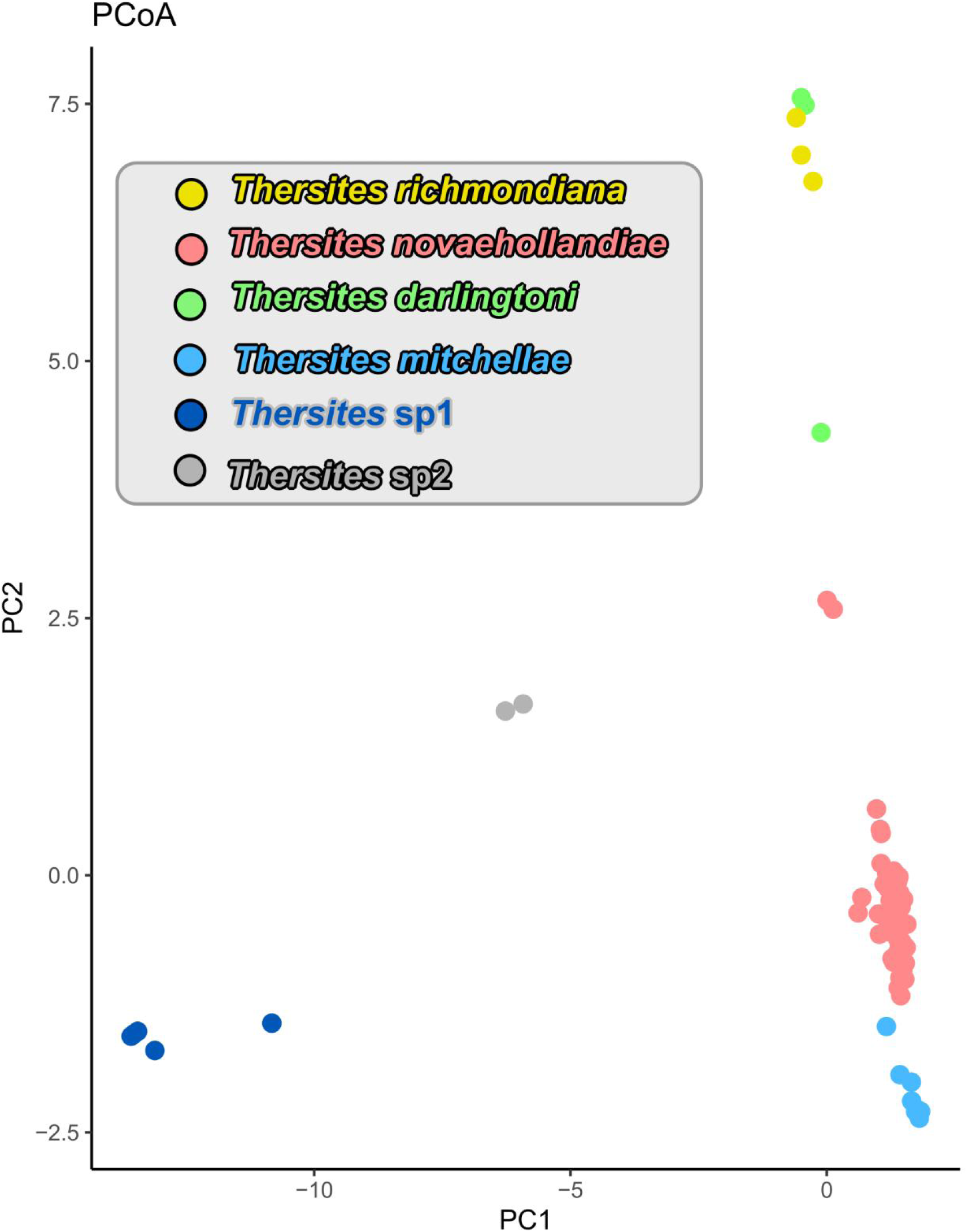
Scatter plots of Principal Coordinate Analysis (PCoA) for DArT sequences

### Molecular phylogenetic analysis

The topologies produced by extended implied weighting (EIW) and implied weighting (IW) under Maximum Parsimony (MP) were identical to one another (Fig. 9A). For K=12, the resulting topology was generally consistent with topologies produced under both EIW and IW for K values between 1 and 100 (CI=0.740, RI=0.940, TL=1018). A single difference in this topology was observed regarding the *T*. *richmondiana* + *T. darlingtoni* lineage. For EIW, the optimal K values, 10, 23, 33, 47, 61-62, 76, 79, 96, and 100 (CI=0.740, RI=0.940, TL=1017), all yielded identical topologies. In contrast, the optimal K values for IW (CI = 0.741, RI = 0.905, TL = 1016) produced two distinct topologies: one group corresponded to the same K values identified as optimal K under EIW, while the other group comprised optimal K values of 52 and 88. Importantly, the strict consensus of these two IW topologies was identical to the topology obtained with the optimal K values under EIW. Applying he equal weighting (EW) method tended to resolve some lineages as polytomies (Fig. S6A). However, this did not affect the phylogenetic position of any nominal taxon. Moreover, compared with the topologies obtained by employing Bayesian Inference (BI) and ML, the EIW/IW approach under MP more frequently resulted in unresolved nodes (Fig. 9B). The topologies of the trees inferred by BI and ML were identical.

**Fig. 9.**
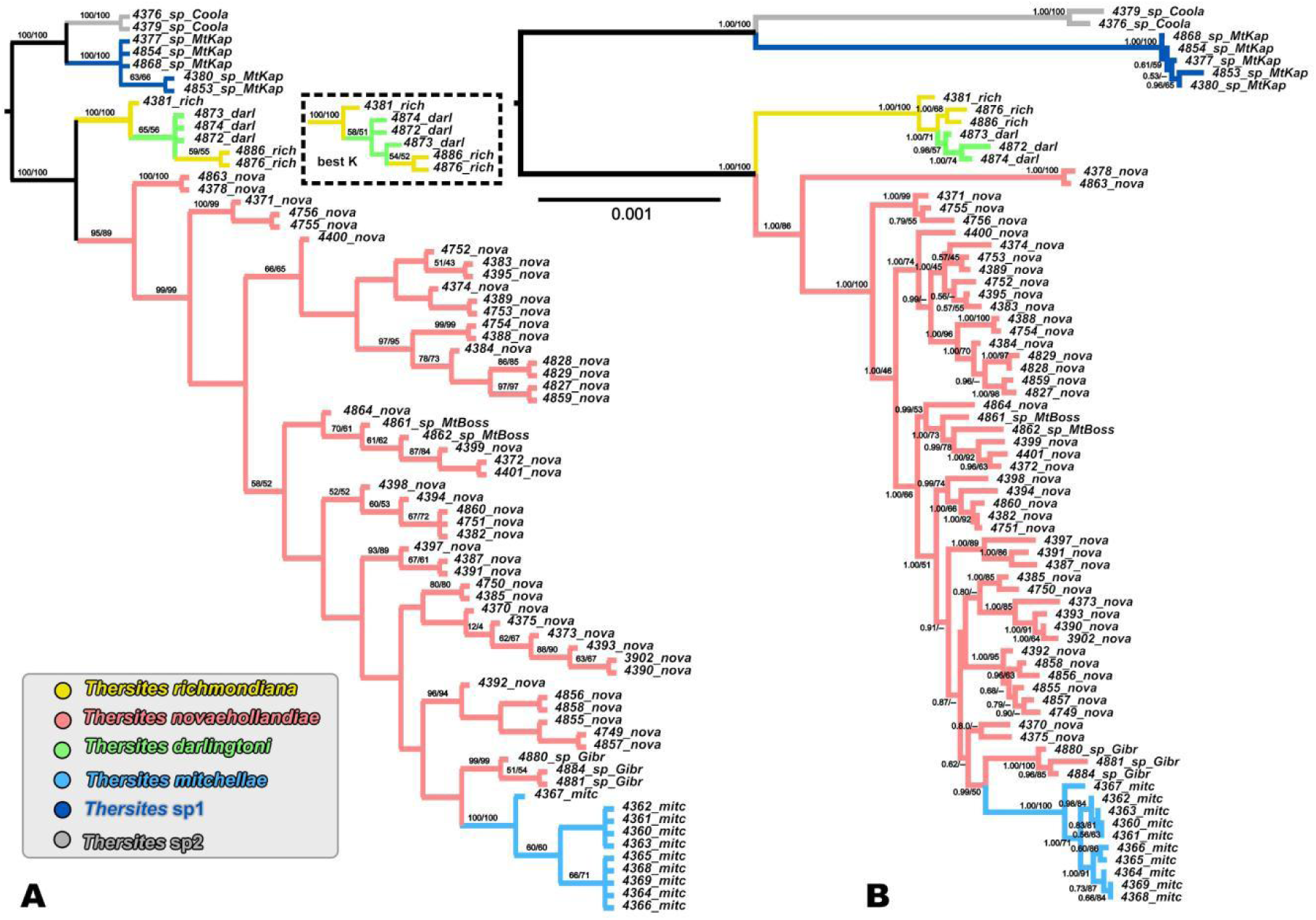
Phylogenetic analyses. A. MP tree under EIW or IW for K = 12 and best K. Numbers on branches indicate support by Jackknifing and symmetric resampling. B. BI tree. Numbers on branches indicating support by Bayesian posterior probabilities and ML bootstraps.

All analyses revealed two principal clades. The first clade comprised *T. richmondiana*, *T. darlingtoni*, *T. novaehollandiae*, and *T. mitchellae*, whereas the second consisted of *T.* sp1 and *T.* sp2. Within the first clade, *T. richmondiana* and *T. darlingtoni* formed a single subclade and *T. novaehollandiae* and *T. mitchellae* formed another.

The clade containing *T. richmondiana*, *T. darlingtoni*, *T. novaehollandiae*, and *T. mitchellae* was maximally supported in all analyses (Parsimony jackknifing, PJK = 100; symmetric resampling, SR = 100; Bayesian posterior probability, BPP = 1.00; bootstrap, BS = 100; ML jackknifing, LJK=100). Within this clade, *T. novaehollandiae* is paraphyletic because *T. mitchellae* occupied a crown group position nested among *T. novaehollandiae* lineages. The monophyly of *T. mitchellae* was maximally supported by jackknifing (PJK = 100) and symmetric resampling (SR = 100) in the MP trees, and by Bayesian posterior probability (BPP = 1.00) and ML bootstrap (BS = 100) and jackknifing (LJK=100) in BI and ML analyses. Furthermore, *T. mitchellae* and *T. novaehollandiae* together consistently formed a strongly supported monophyletic group (PJK = 95, BPP = 1.00, SR = 89, BS = 86, LJK=100). *Thersites richmondiana* and *T. darlingtoni* clustered together with maximal support (PJK = 100; SR = 100, BPP = 1.00; BS = 100, LJK=100) in all analyses. However, *T. darlingtoni* occupied the crown position, rendering *T. richmondiana* paraphyletic. The monophyly of *T. darlingtoni* was strongly supported by BI (BPP=0.98) and likelihood jackknifing (LJK=100) but weakly by ML bootstrapping (BS=74).

The second main clade comprising *T*. sp1 and *T*. sp2 is maximally supported across all analyses (PJK = 100; SR = 100; BPP = 1.00; BS = 100, LJK=100), with both taxa forming a maximally supported sister group (PJK = 100; SR = 100; BPP = 1.00; BS = 100, LJK=100).

The phylogenetic network (Fig. 10) indicates that *Thersites mitchellae* is nested within *T. novaehollandiae*, forming a network. *T. darlingtoni* and *T. richmondia* form a lineage, but both are polyphyletic. *T. sp1* and *T. sp2* are monophyletic and are sister taxa to each other.

**Fig. 10.**
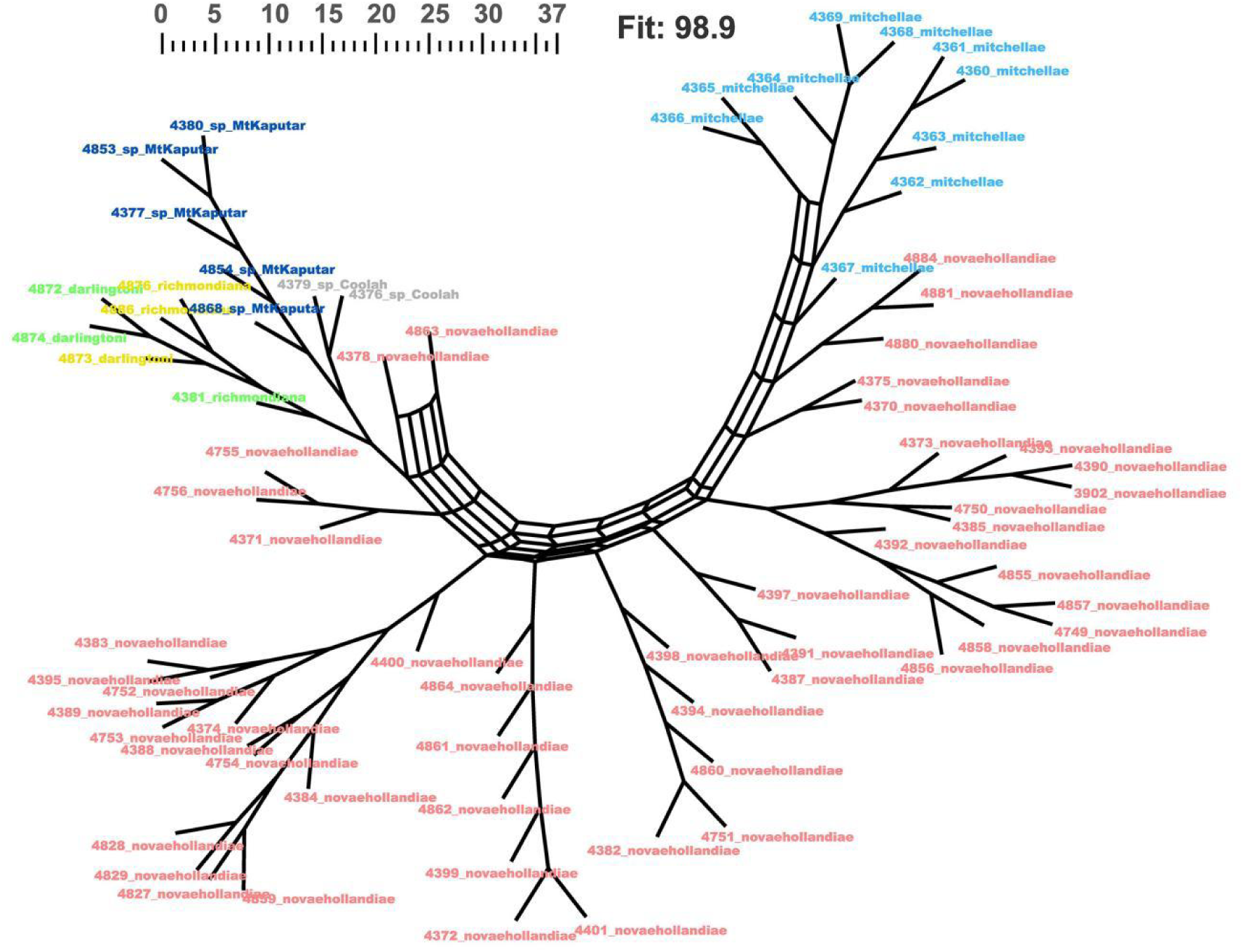
Phylogenetic network generated using the consensus outline method, summarizing the results from parsimony-based phylogenetic networks.

### Morphological character optimization analysis

The continuous character optimization (CCO) for shell dimensions and size of reproductive organs (Fig. S7) revealed no apomorphic characters. Most characters exhibited homoplasy at species level. No character changes were observed at node *T. mitchellae* + *T. novaehollandiae*. The lineage of *T. darlingtoni* + *T. richmondiana* was characterized by shell width and height, as well as the lengths of the phallus, epiphallus2, and flagellum. *Thersites* sp1 was characterized by shell height and width, while *Thersites* sp2 was characterized by the lengths of epiphallus2 and flagellum. No homoplastic changes originated at the nodes of *T.* sp1 + *T.* sp2 or in the other four described species.

Heatmaps mapped on to the phylogenies (Fig. 11) indicated that continuous characters varied substantially within and among species. Indeed, we found no clear differences between the inter- and intraspecific ranges of variation. However, we found that *T*. sp1 and *T*. sp2 were consistently smaller than all other species. *Thersites mitchellae* and *T. novaehollandiae* were highly variable while *T. darlingtoni* differed little from *T. richmondiana*. The Kruskal-Wallis test heatmaps plotted onto the phylogeny (Fig. 14) revealed statistically significant differences between the species pair *T*. sp1 + *T*. sp2 and the group of *T. darlingtoni*, *T. richmondiana*, *T. mitchellae*, and *T. novaehollandiae* in shell width and height as well as lengths of phallus, epiphallus1, epiphallus2, and flagellum. In addition, significant differences were observed between *T*. sp1 and *T*. sp2 in shell width and height as well as in the lengths of epiphallus2 and flagellum. Comparisons of *T. darlingtoni* and *T. richmondiana* with *T. mitchellae* and *T. novaehollandiae* revealed significant differences in shell width and height, epiphallus2 and flagellum. The differences between *T. mitchellae* and the crown group of *T. novaehollandiae* were significant for shell width and the lengths of phallus and flagellum. Other significant differences have also been detected among various *T. novaehollandiae* lineages.

**Fig. 11.**
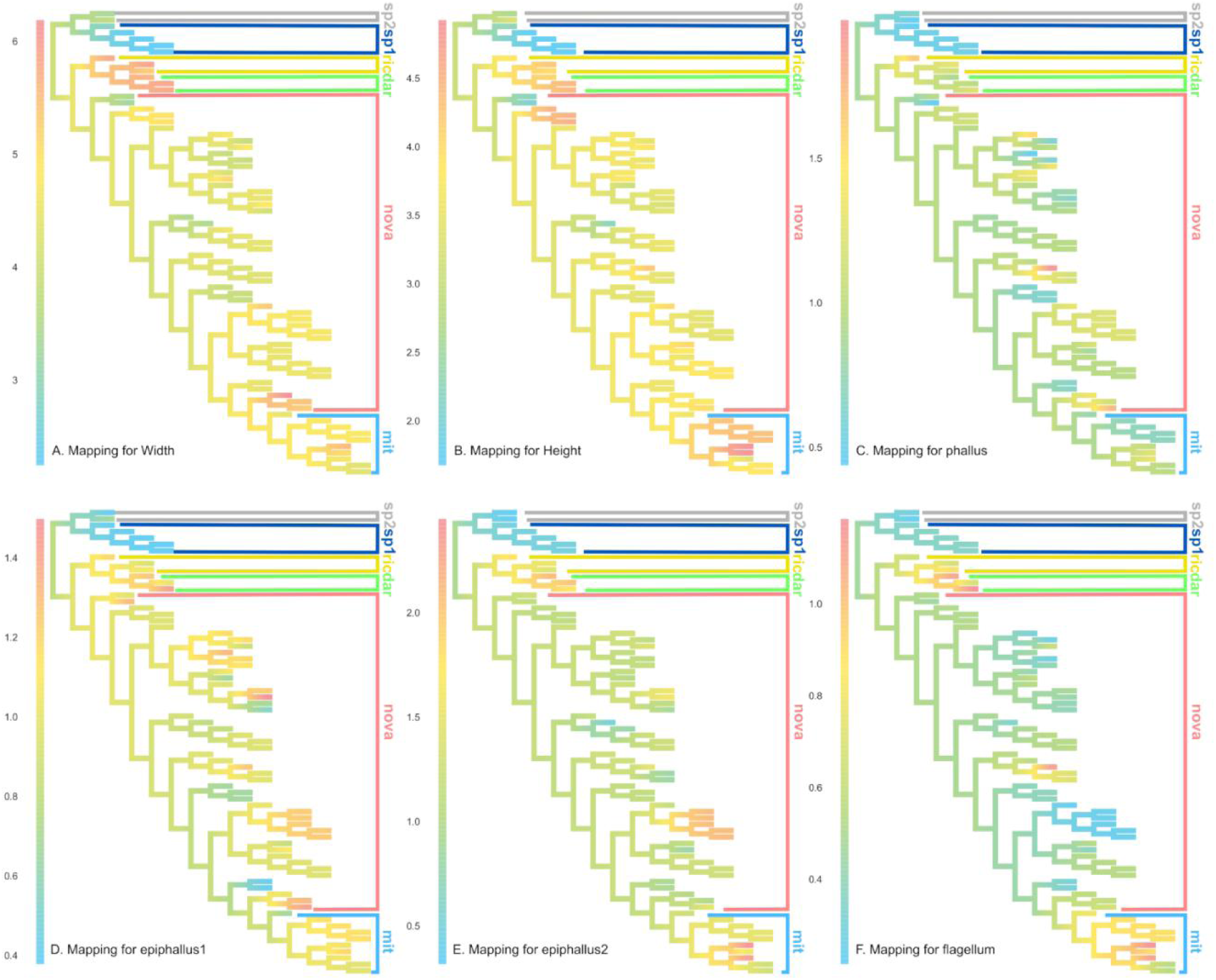
Continuous characters changes mapped on the BI tree.

The CCO for landmarks suggested that nearly all nodes have apomorphic characters (Fig. 12). The thin plate splines suggested that aperture semi-landmarks varied substantially among the phylogenetic trees, even at the individual level. The heavily varied body whorl semi-landmarks and landmarks were observed at the nodes of *Thersistes* sp1 + *T*. sp2, *T*. sp1, *T. richmondiana* + *T. darlingtoni* and some specific *T. novaehollandiae* lineages.

**Fig. 12.**
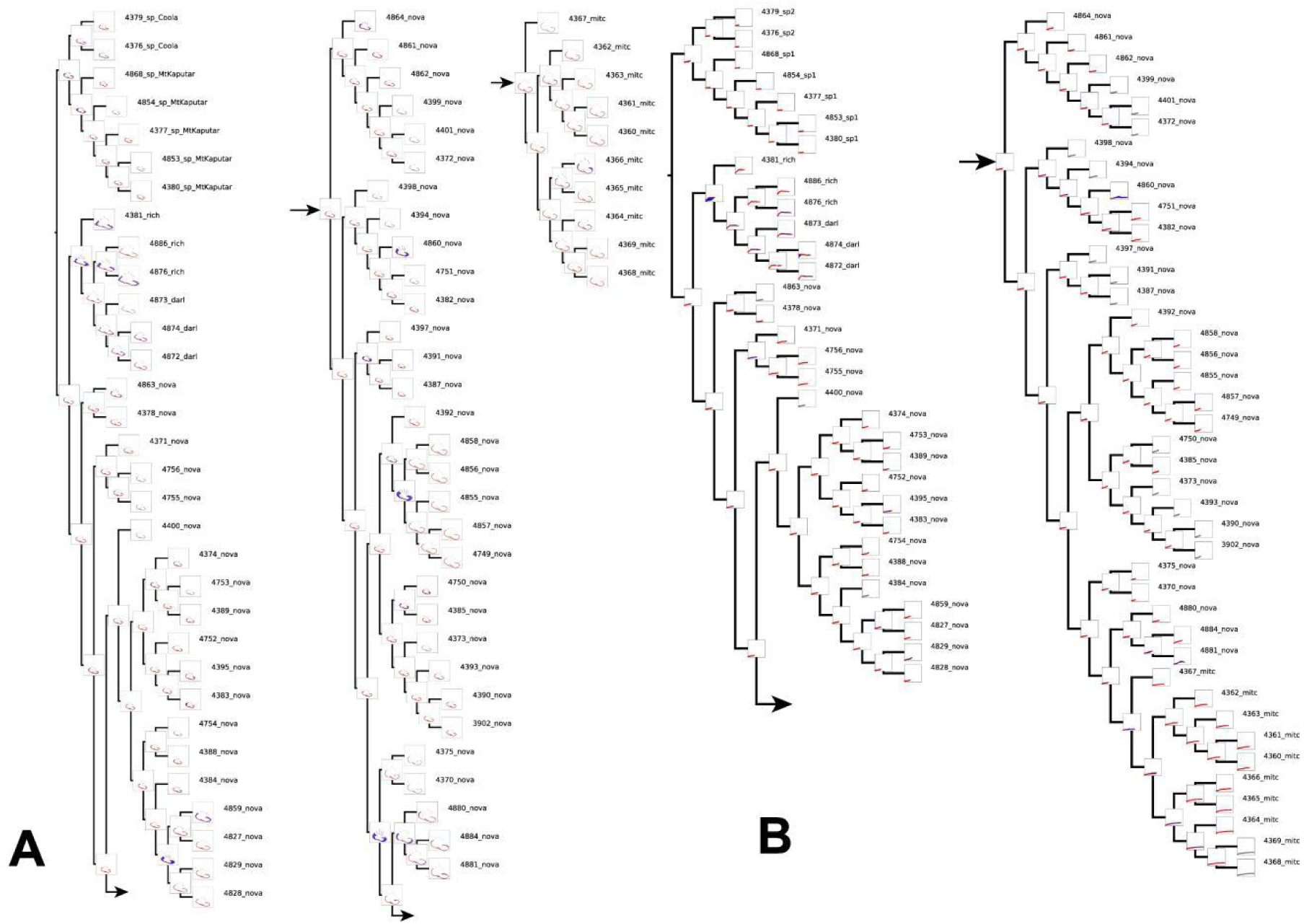
Landmarks optimization for shells (A) and 50% flagellum (B). Blue lines indicate apomorphies and homoplasic characters.

For the ancestral node of *T*. sp1 + sp2, a depressed shell was identified as an apomorphic character state. The shell of *T*. sp1 is even more depressed than the ancestral character state at this node. The lineage of *T. darlingtoni* + *T. richmondiana* has a keeled shell as an ancestral character state with a higher spirea nd a blunt flagellum tip. For the lineage of *T. mitchellae* + *T. novaehollandiae*, we identified no landmark apomorphies. Only *T. mitchellae* and its sister clade had a globose body whorl and a high spire as ancestral state.

Our reconstructions suggested that some lineages of *T. novaehollandiae* underwent similar changes in shell shapes with the body whorl being either more depressed or more globose.

Silhouette scores for all landmarks (Fig. 13) indicated that the optimal clustering was achieved with two clusters. But these clusters were not fully consistent with the taxonomic null hypothesis. Based on flagellum shape, *T. novaehollandiae*, *T. darlingtoni*, and *T. richmondiana* are distributed across both clusters, whereas *T.* sp1, *T.* sp2, and *T. mitchellae* occurred exclusively in cluster 1. When considering the seven shell landmarks, all species were represented in both clusters except *T.* sp1 and *T.* sp2.

**Fig. 13.**
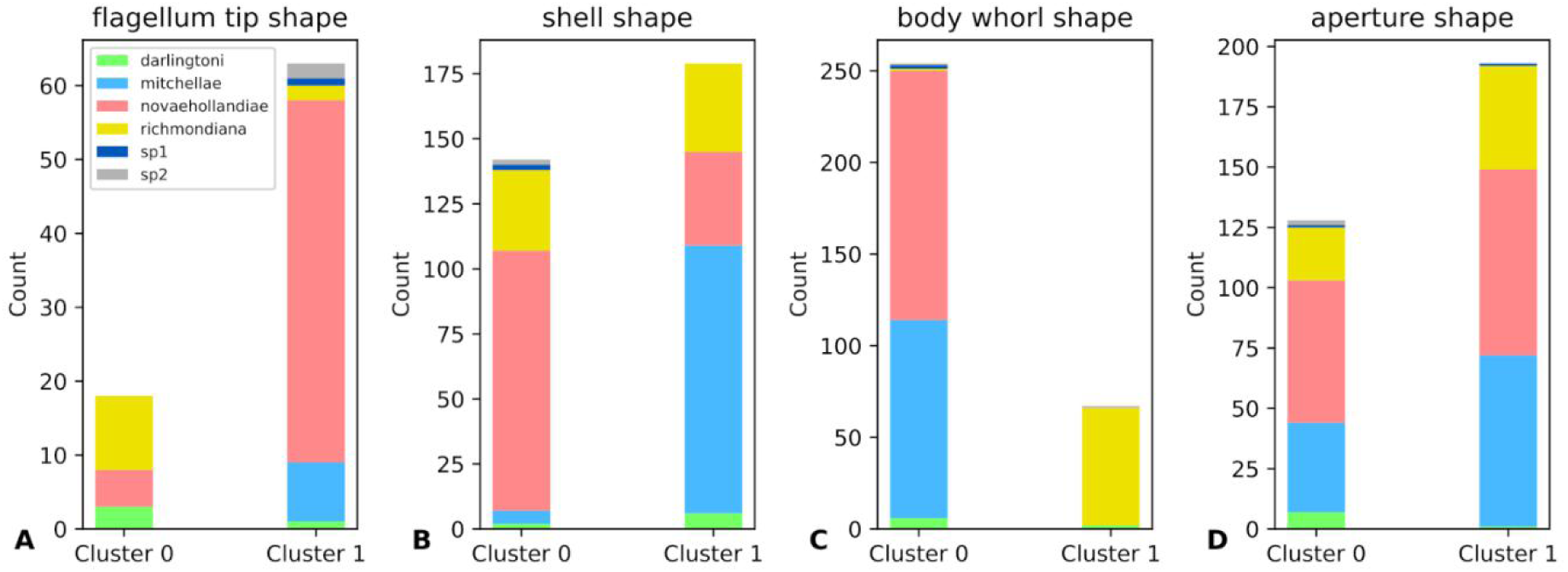
Bar charts of K-means clustering for landmarks and semi-landmarks. The flagellum tip shape represents 50% of flagellum length landmarks. Shell, body whorl and aperture shapes separately represent the landmarks, semi-landmark curve 1 and semi-landmark curve 2.

Analysis of body whorl semi-landmarks showed that *T.* sp2 was present in both clusters, nearly all *T. richmondiana* were confined to cluster 1, and almost all *T. darlingtoni* were found in cluster 0, with the remaining species occurring in a single cluster. Regarding aperture shape, *T.* sp2 appeared only in one cluster, nearly all *T. darlingtoni* grouped in a single cluster, while the other species were distributed across both clusters.

Under both fast and slow optimization (Fig. 14), the only apomorphy that has been identified supports the sister group relationship between *T.* sp1 and *T.* sp2. This apomorphy is an open umbilicus. By contrast, we identified multiple homoplasies across various nodes in the tree.

**Fig. 14.**
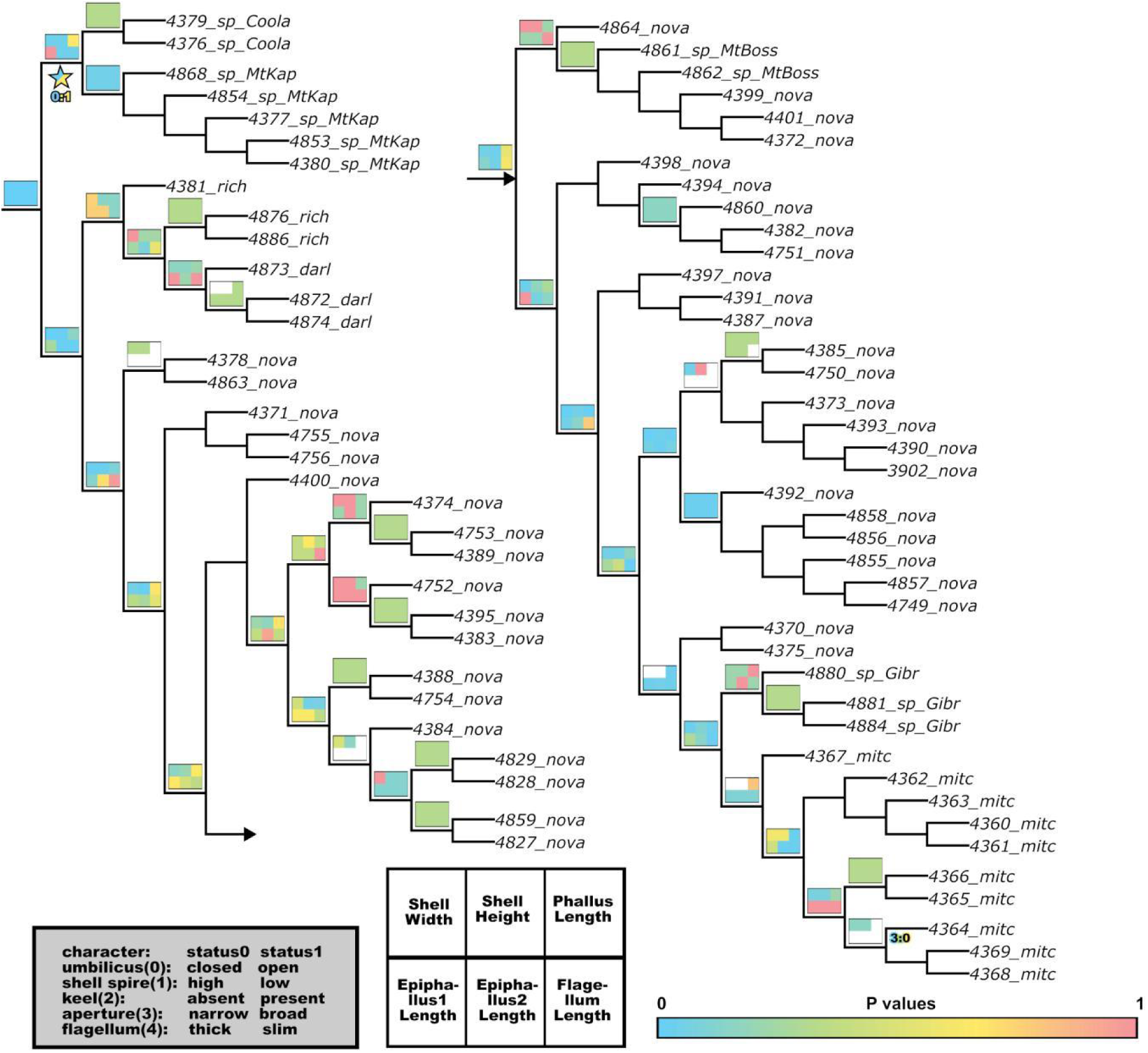
Kruskal-Wallis test heatmaps of character measurements are marked beyond branches of the phylogeny. The homoplasic and apomorphic characters under fast, slow, and unambiguous character optimization are marked under the branches of phylogeny. Six-box heatmaps show the p-values calculated for measurements of shell width and height, measurements of phallus, epiphallus1, epiphallus2, and flagellum. These p-values are derived by comparing the branch’s immediate child nodes to all its descendant nodes. Apomorphic characters are marked using stars. Characters are labeled in the ‘character number: stat number’ format. Discrete characters (characters 1-4) were derived from landmarks categorized using K-means clustering, along with one qualitative character (character 0, indicating a open or closed umbilicus). Only open umbilicus under fast/slow character optimization is the apomorphic character for *Thersites* sp1 and sp2.

**Fig. 15.**
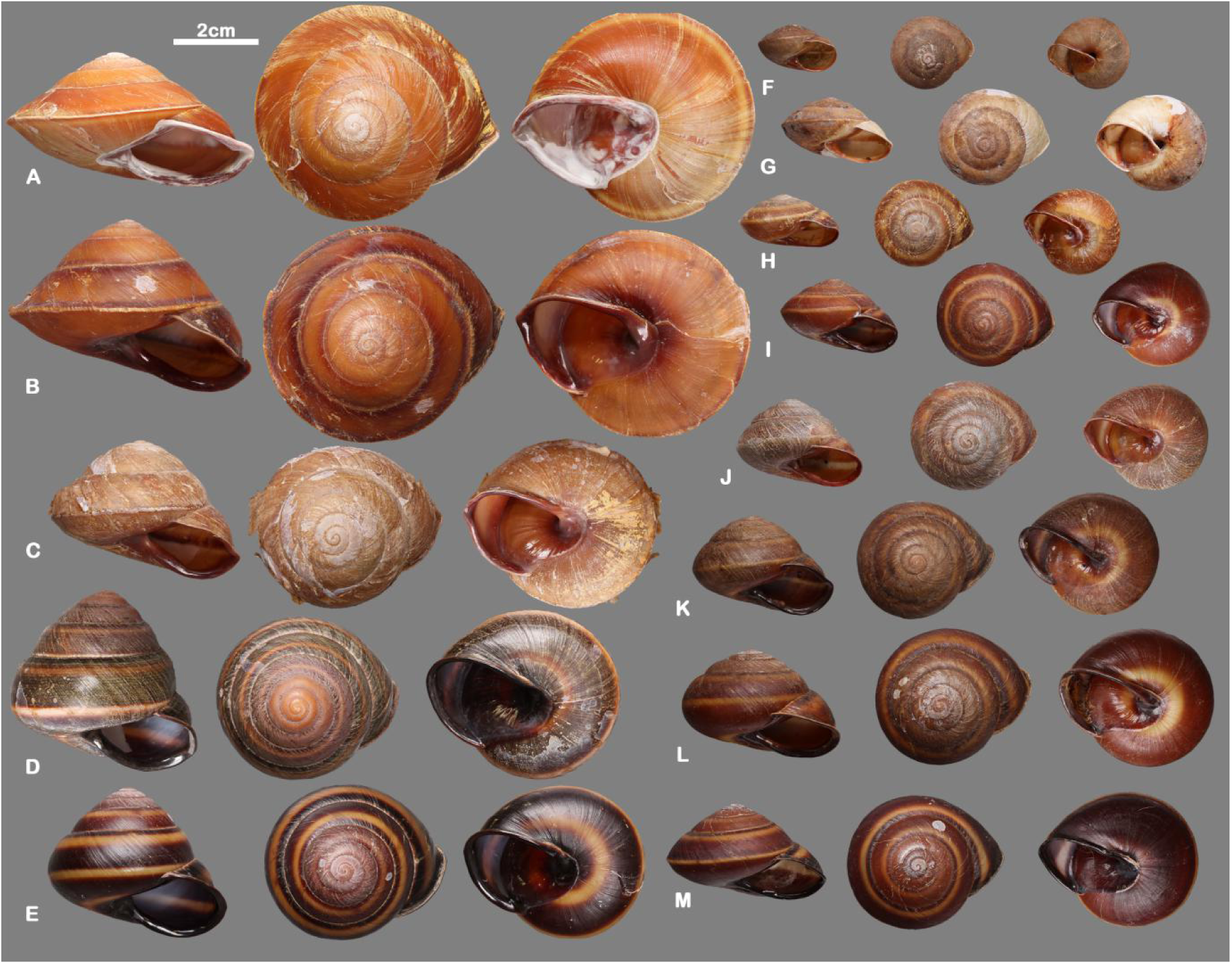
Shells of specimens. A—B *Thersites richmondiana*, QM MO4235, QM MO25801spec2; C *T. darlingtoni*, QM MO59910; D—E *T. mitchellae*, AM C. 584219, AM C. 584209; F *T. kaputarensis*, AM C.615708; G *T. coolahensis*, AM C.2721928; H —M *T. novaehollandiae*, QM MO36956spec1, AM C. 582443, QM MO 66656, AM C. 272195, AM C. 575474, AM C. 584422.

**Fig. 16.**
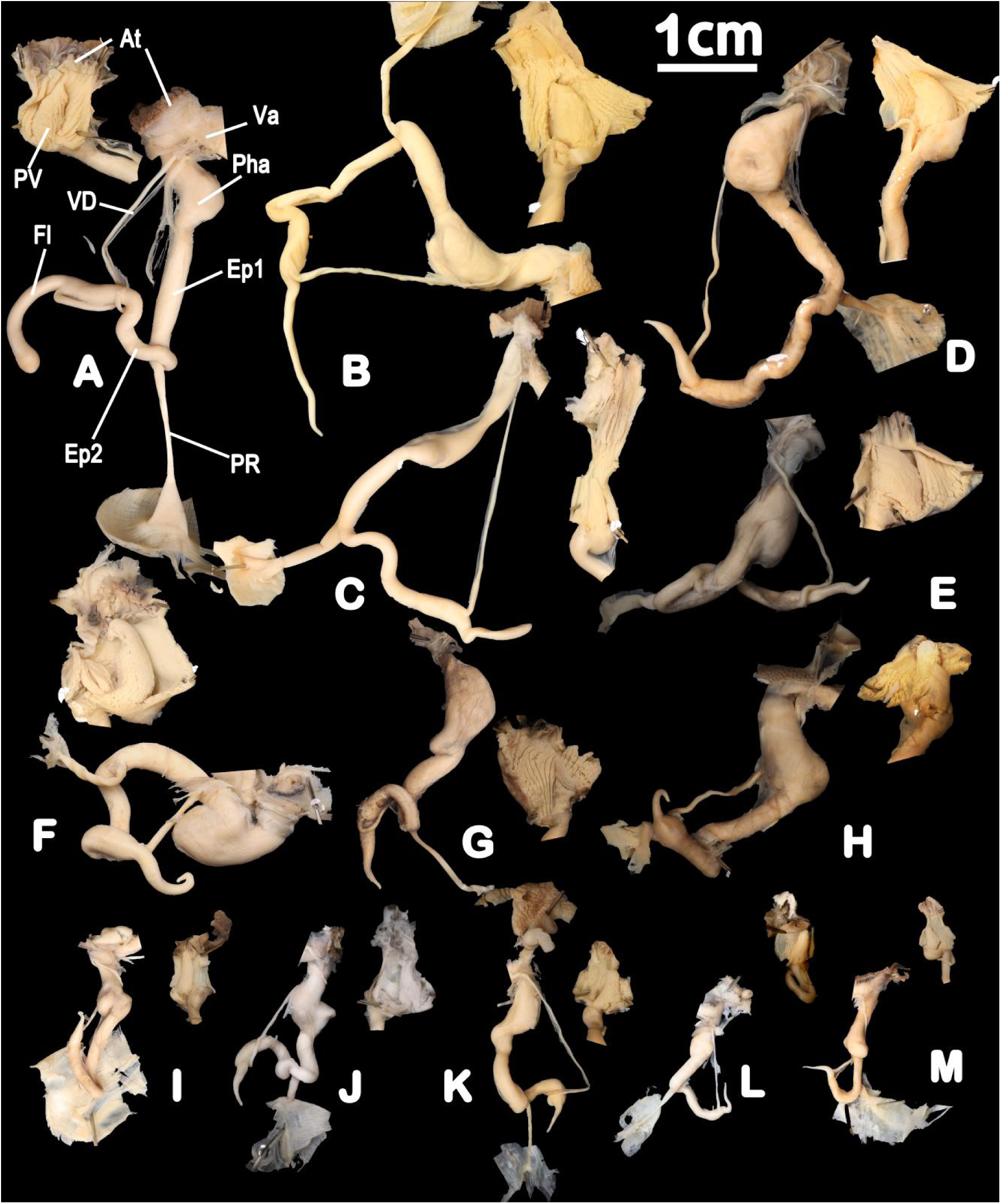
Male genital organs and opening of atrium showing penial verge and penial pilasters. A *Thersites darlingtoni*, AM C.59910spec1; B—C *T. richmondiana*, QM MO4235, QM MO25801spec2; E—F *T. mitchellae*, AM C. 584209, AM C. 584219; D,F—K *T. novaehollandiae*, QM MO17029spec3, AM C. 272195, AM C. 575474, AM C. 582443, AM C. 584422, AM C. 36956spec1; L *T. kaputarensis*, AM C.615708; M *T. coolahensis*, AM C.2721928. At—atrium; Ep1—epiphallus1; Ep2—epiphallus2; Fl—flagellum; Pha—phallus; PR— penial retractor; PV—penial verge; Va—vagina; VD— vas deferens.

## Discussion

### Taxonomic significance of shell and genital morphometrics

In the latest taxonomic revision, Bishop (1978) employed simple ratios in shell and genital dimensions to distinguish *Thersites* species claiming that this can be achieved by means of two independent, continuous character states, the shell height/width ratio and the ratio between flagellum and proximal epiphallus length. Bishop (1978) stated that *T. mitchellae* exhibited a height–width ratio greater than 0.8, whereas *T. richmondiana*, *T. darlingtoni* and *T. novaehollandiae* supposedly have ratios smaller than 0.7. In addition, Bishop (1978) stated that the ratio between flagellum length and length of the proximal epiphallus was lower than 0.25 in *T. novaehollandiae*, about 0.5 in *T. richmondiana* and *T. darlingtoni*, and about 1.0 in *T. mitchellae*. However, when giving these ratios, Bishop (1978) did not clearly define what exactly the ‘proximal part of the epiphallus’ was supposed to be. Independent of this detail, we found that none of these stated ratios were actually diagnostic. In fact, we found that no species was clearly defined by any of these two ratios for species overlapped with one another in these morphometric parameters (Figs S2, S3).

While Bishop (1978) synonymized *T. darlingtoni* with *T. richmondiana*, Stanisic et al. (2010) treated both as accepted species stating that *T. darlingtoni* can be distinguished from *T. richmondiana* by smaller shell size and by having a rounded instead of keeled shell periphery. However, this statement was not backed up by an analysis of any data (Figs 4-6).

We tested the classification last proposed by Stanisic et al. (2010) by rigorously analysing the supposedly diagnostic character states in a statistical framework as well as conducting genomic analyses. Our comparative analyses of morphological traits revealed that the four currently accepted *Thersites* species cannot be reliably distinguished using shell shape characteristics nor by using reproductive traits, such as flagellum length. In fact, based on the character distributions alone, we cannot recognize any clearly distinguishable group. The Shapiro–Wilk test indicated that shell dimensions, such as shell width and shell height, and flagellum length, were not normally distributed. However, histograms failed to reveal distinct states in any of these characters that could be used to distinguish taxa. Moreover, the measurements of genital organs and shells revealed no clear separation between the amounts of intraspecific and interspecific variation in the taxa as previously delineated (Figs 4, S1).

### Species monophyly

As the current morphology-based classification has not withstood statistical scrutiny, but was also not rejected outright, we also employed a molecular phylogeny to put the initial taxonomic hypothesis to the test. The phylospecies concept defines species as the smallest monophyletic group that is distinguished from other such groups by one or more apomorphies (Wilkins, 2018). Consequently, it is also required that species are monophyletic in phylogenetic trees. Our molecular phylogeny of *Thersites* based on analyses of nuclear loci revealed that two species were non-monophyletic, however. First, we showed that *T. novaehollandiae* encompassed several lineages, one of which is represented by *T. mitchellae* rendering *T. novaehollandiae* paraphyletic. This finding corresponds well with the mitochondrial phylogeny of Hugall and Stanisic (2011), who also recovered *T. novaehollandiae* as paraphyletic with respect to *T. mitchellae*. This finding is consistent with the phylogenetic network analyses, in which *T. novaehollandiae* and *T. mitchellae* are nested together. Second, we also found that *T. darlingtoni* and *T. richmondiana* were mutually non-monophyletic (Figs 9, S6). Accordingly, neither *T. mitchellae* nor *T. darlingtoni* as currently delimited should be accepted as distinct species based on the criteria of monophyly in combination with the rules regarding nomenclatural priority. However, before a final taxonomic conclusion can be drawn from the available tree, we need to test if species can be delineated in alternative ways, to form monophyletic groups that are supported by apomorphies. To this end, we have embarked on a strategy of character optimization as outlined in the following.

### Morphological character optimization

We use character optimization, i.e., the mapping of observed character states at the tree tips and the reconstruction of ancestral character states across the given tree, to identify potential morphological apomorphies at the nodes across the tree topology. By doing that, we simultaneously test any possible alternative taxonomic hypothesis through the mapping of both discrete and continuous morphological characters while seeking to identify apomorphies in the reconstructed ancestral character states.

Discrete characters can be and have been readily used to identify apomorphies. However, in the present case the number of discrete characters is insufficient even to support the revised taxonomic hypothesis that there are just two accepted species, *T. novaehollandiae* and *T. richmondiana*, and two undescribed species, *Thersites* sp1 and sp2. When ignoring the missing data, there is a single character with two states (umbilicus open/closed) that distinguishes the two undescribed species from the other species as a synapomorphy (= open umbilicus).

In search of additional apomorphies, we categorized the landmark datasets for shell and genital features. This has not been done before, although there are some methods to use landmarks in the framework of cladistic analyses (e.g. Goloboff and Catalano, 2016). Our Silhouette score analyses suggested that the most robust categorizations for these landmark datasets always yielded two states, with individual species often displaying both. Consequently, our attempt to derive apomorphies by discretizing the landmark covariance matrix failed because of low homology in the shapes described, echoing earlier efforts by Zelditch et al. (1995).

Our CCO for landmarks suggested that the ancestral morphotype of (*T.* sp1 + *T.* sp2) had a small, depressed shell. Moreover, we found that *T.* sp1 has a more depressed shell than *T.* sp2, and that *T. richmondiana* has a stronger keel and higher spire than *T. novaehollandiae*. *T*hersites *richmondiana* has a blunt flagellum tip while *T. novaehollandiae* has not. CCO for landmarks can reveal substantial shape changes at ancestral nodes that may provide useful information for species delimitation. However, in the absence of clearly defined boundaries for shape variation limits the predictive value of this approach. We conclude that CCO for landmarks has an advantage to categorising landmark characters because it provides not just a summary of observations and induction; it also allows for deduction. Just as what Schleiden (1842) said “a collection of naked facts is still far from being science, just as mere building material is not yet a temple”. This method also maintains the original continuous character, doesn’t lose any information.

Other methods, such as Contrasting Composite and Reductive Coding (Torres-Montúfar et al., 2018), have been proposed to resolve homoplasy issues that represent interpretation errors of characters (Nixon and Carpenter, 2002). As noted by Torres-Montúfar et al. (2018), complex characters should be iteratively decomposed into simpler, more homologous units. However, when the traits (such as shell shape) are heavily influenced by environmental factors (Goodfriend, 1986), this process becomes particularly challenging.

The CCO for shell dimensions and size of reproductive organs revealed that homoplastic characters have been present at all nodes of the phylogenies. Therefore, we cannot readily identify species boundaries on the tree topologies by using this method. Statistical tests did also not recognize natural groups when applied globally to all samples (Figs 4, S1). However, the Krustal-Wallis tests applied at specific nodes of the trees were suitable to identify clades with distinct morphological character states. In our phylogeny, three nodes supported the four morphologically distinct lineages according to the Krustal-Wallis tests: First, the sister groups (*T.* sp1 + *T.* sp2) and (*T. novaehollandiae* + *T. richmondiana)* differed from one another in the size of all organs (i.e., shell width and height, lengths of phallus, epiphallus1, epiphallus2, and flagellum; p < 0.05). Second, the species in each of these two sister pairs could be distinguished from one another by means of shell width and height as well as in the length of epihphallus2 and flagellum (p < 0.05). This result is consistent with the outcomes of CCO for shell dimensions and size of reproductive organs. We found that the following species are diagnosed by these diagnostic character states: *Thersites* sp1 (lengths of epiphallus2 and flagellum), *T*. sp2 (lengths of epiphallus2 and flagellum), *T. richmondiana* (shell width and height, lengths of epihphallus2 and flagellum). By contrast, *T. novaehollandiae* has no continuous character support. Compared to directly mapping continuous characters onto phylogenetics, the Kruskal-Wallis test is able to identify the boundaries of continuous character variation.

It is critically important to recognize that statistical clustering primarily reflects similarity rather than true homology (Sneath and Sokal, 1973). We try to avoid this caveat here by applying statistical tests only to natural groups that have been recognized in a phylogenetic framework. While across the entire group, most morphological characters overlap in their ranges, we found that they differ with statistical significance at specific nodes in the phylogeny. We argue that these morphological differences are apomorphies, especially when they were also identified by continuous homoplastic character states.

Finally, our findings challenge the gen–morph species concept (Hong, 2020), which requires species to possess at least two statistically discontinuous apomorphies. This criterion is unlikely to be met in groups that are characterized by few, overlapping characters. In a saturated morphospace (Budd, 2021), morphological variation tends to be continuous, reflecting gradual shifts rather than clear-cut, discrete differences (Foote, 1994; Paup, 1966; Erwin, 2006). Such gradual shifts are also encountered in the present case. A strict application of the gen-morph species concept to the case of *Thersites* would lead to the recognition of just a single species as an artifact of lacking characters rather than a realistic reflection of the true diversity of the group. This illustrates that by enforcing a prescribed set of diagnostic criteria, one may only detect a subset of the actual species-level diversity (see also Wiley and Lieberman, 2011). Our results underscore that attempts to integrate all aspects of species delimitation into a single framework are unlikely to be universally applicable, such as the gen-morph species concept. Therefore, we agree in principle with the critique of Mayden’s (1997), that species concepts should not be operational.

### How to use morphological characters

We used *Thersites* as an example to explore how best to incorporate analyses of continuous characters to recognize apomorphies for taxa with a limited number of discrete characters. To this end, we aim to objectively define character status. As Pimentcl & Riggins (1987) and Stevens (1991) pointed out that without delimiting character status objectively there can be no meaningful morphological character analyses (“garbage in, garbage out”). Our study makes steps toward the mathematization of systematics, aligning with the ideal of the ‘*mathesis universalis*’, which Hennig (1949) described as ‘all goals of all true science’. Hence, systematics is not a ‘meeting ground of art and science’ as Davis & Heywood (1963) have claimed.

## Acknowledgements

Thanks go to Alison Miller (AM), Jennifer Caiza (AM), John Stanisic (QM), Darryl Potter (QM), Jennifer Trimble (MCZ), Gonzalo Giribet (MCZ), Jon Ablett (NHM) for making material under their care available to this study and/or for providing photographs of specimens. We thank Craig Stehn for helping with fieldwork. We thank the Willi Hennig Society for providing access to the software TNT. The molecular component of this study was funded by the NSW Department of Climate Change, Energy, the Environment and Water, which is thankfully acknowledged.

## Supplementary File

**Supplementary Fig. 1.**
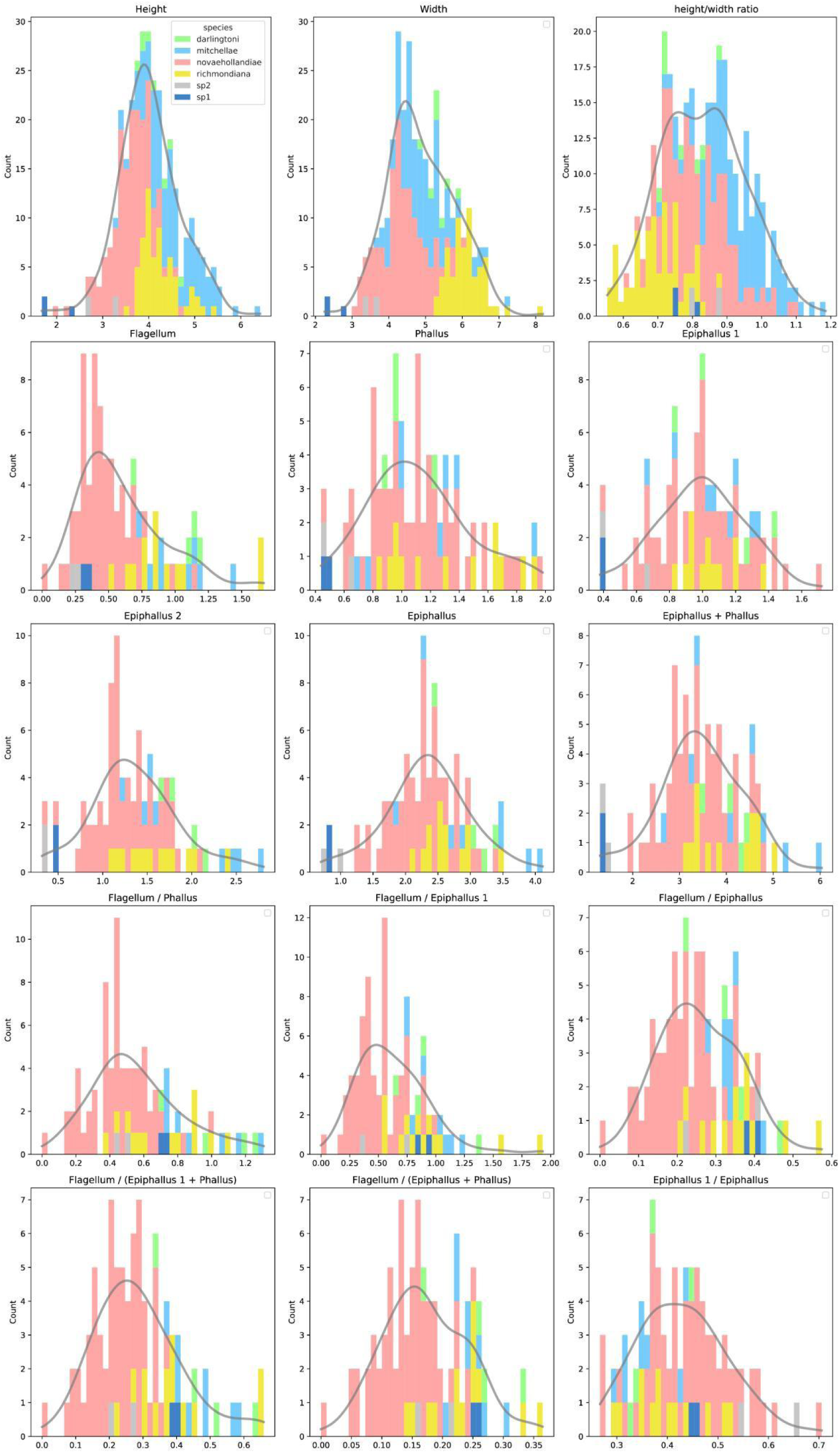
Histograms of shell height, width, height/width ratio and genital measurements and their ratios of *Thersites*. Species.

**Supplementary Fig. 2.**
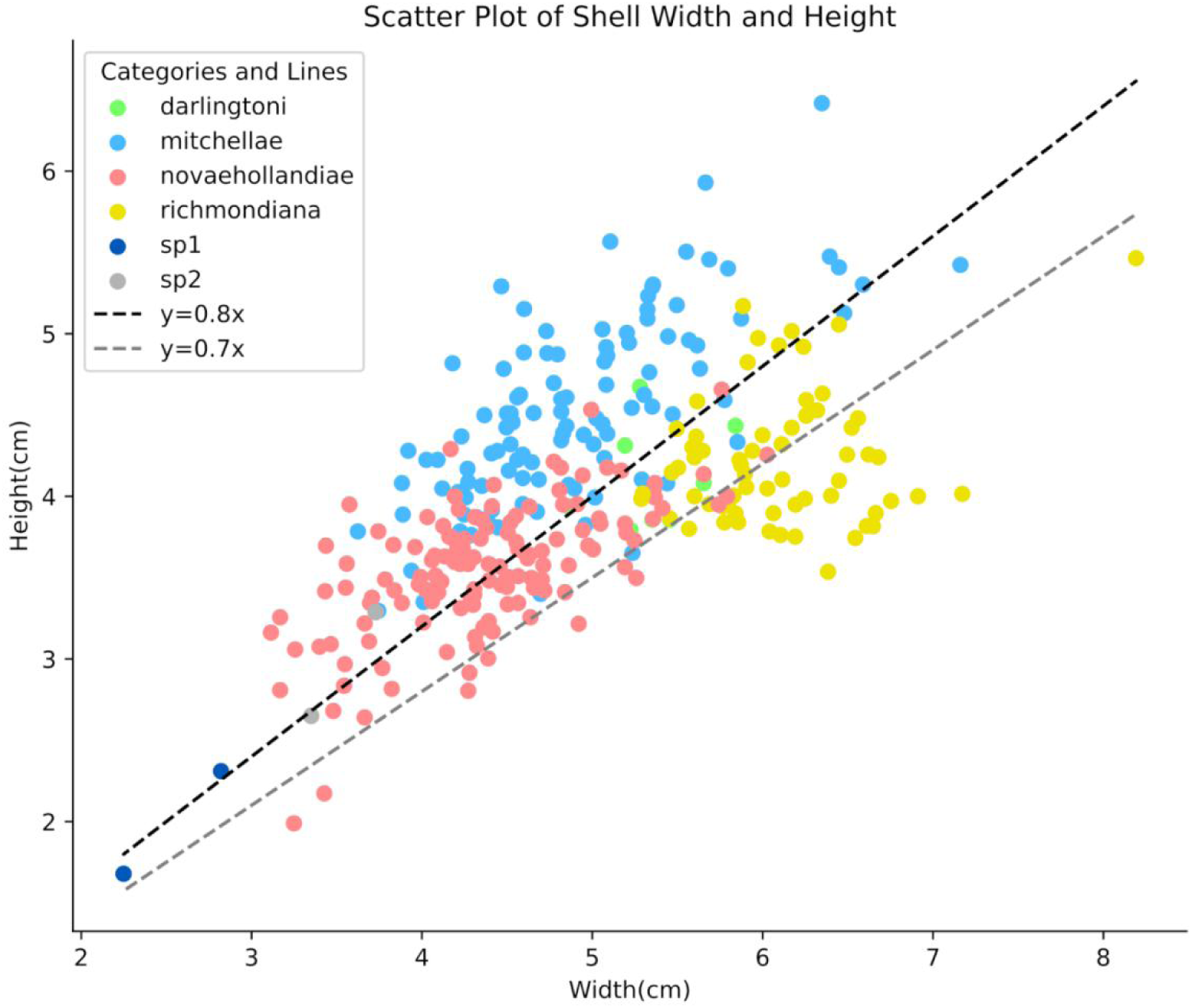
Scatter plots of shell width and height.

**Supplementary Fig. 3.**
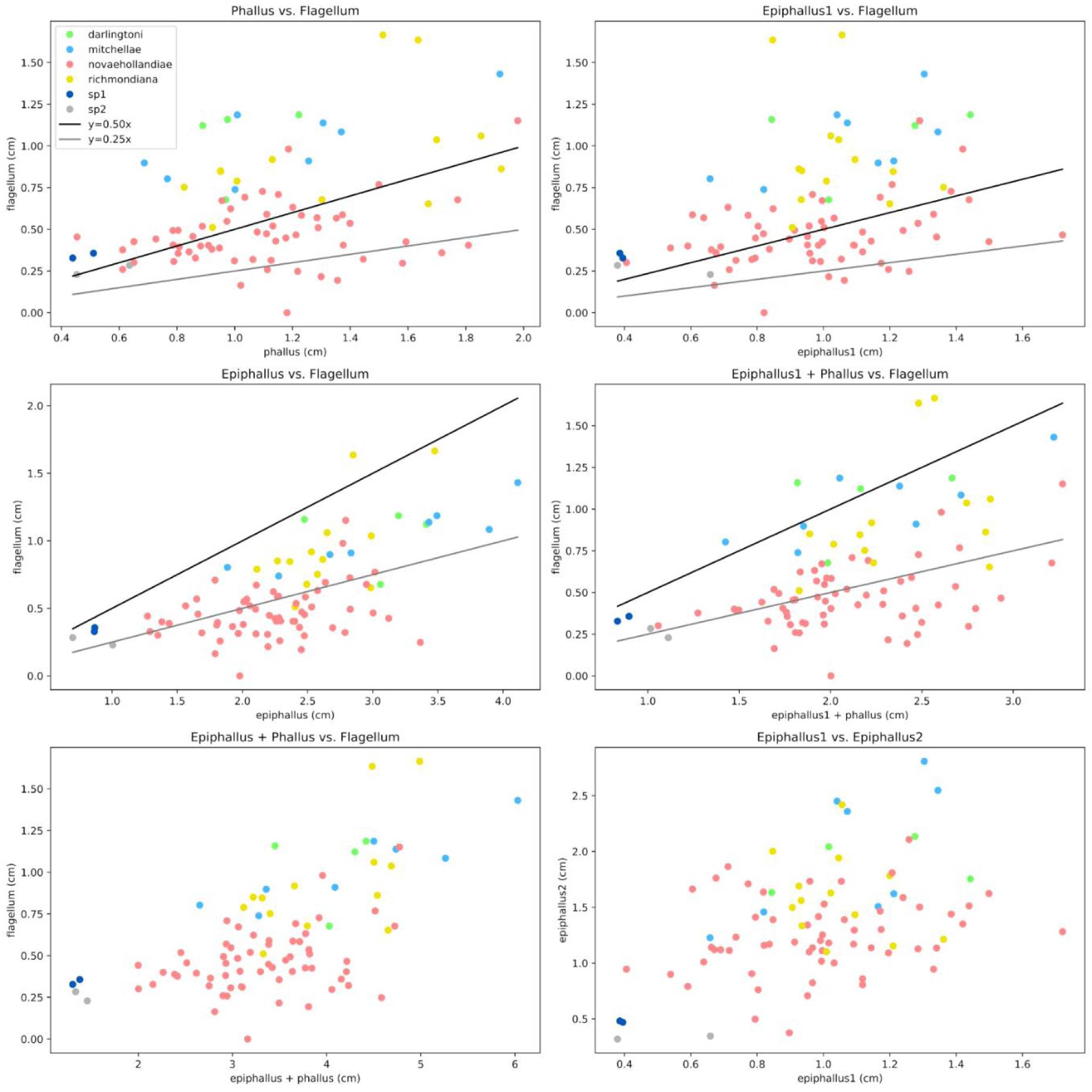
Scatter plots of genital measurements.

**Supplementary Fig. 4.**
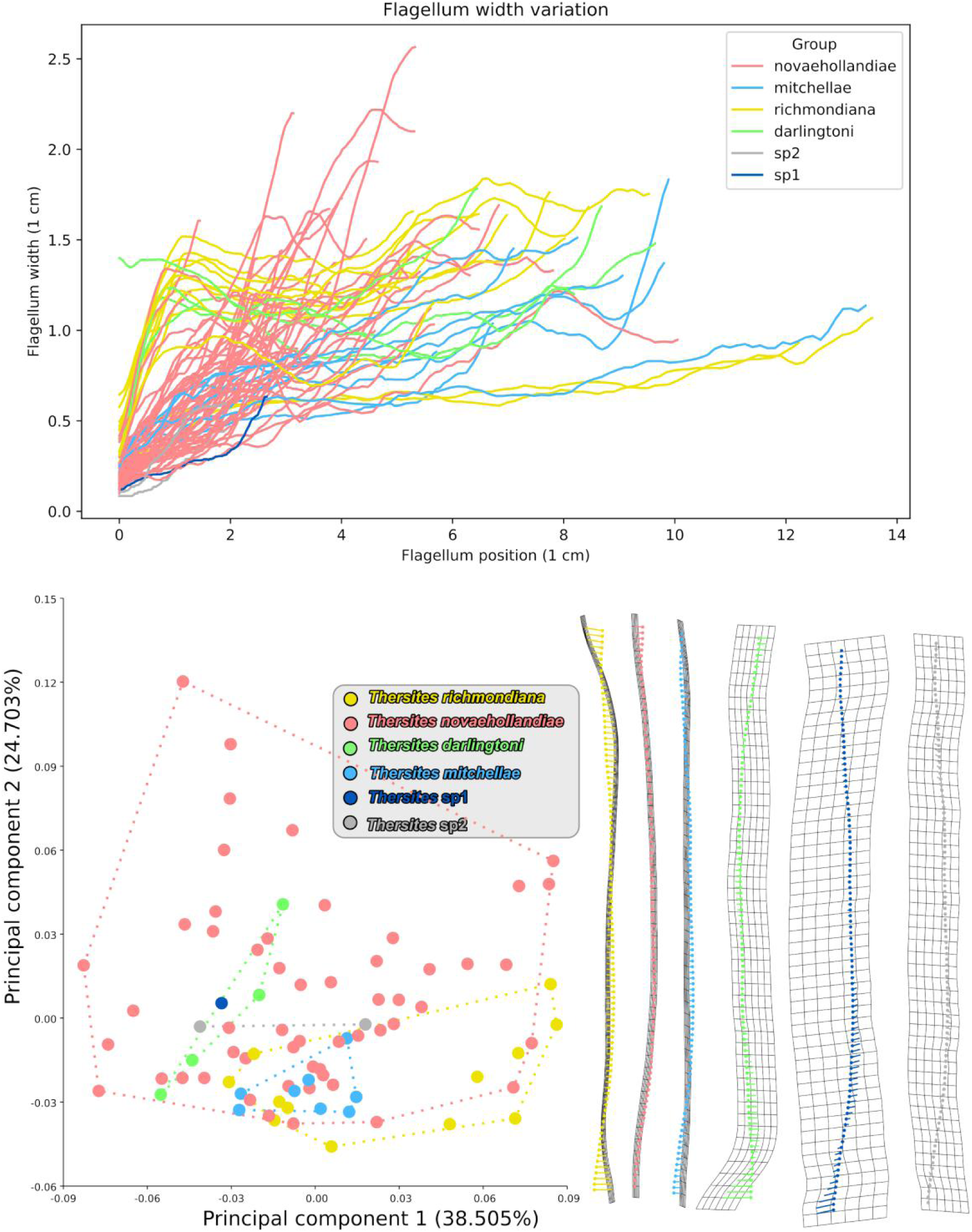
Curves of 100% length flagellum shapes and scatter plots of principal component analysis for 100% length flagellum with thin plate spline grid maps for each species.

**Supplementary Fig. 5.**
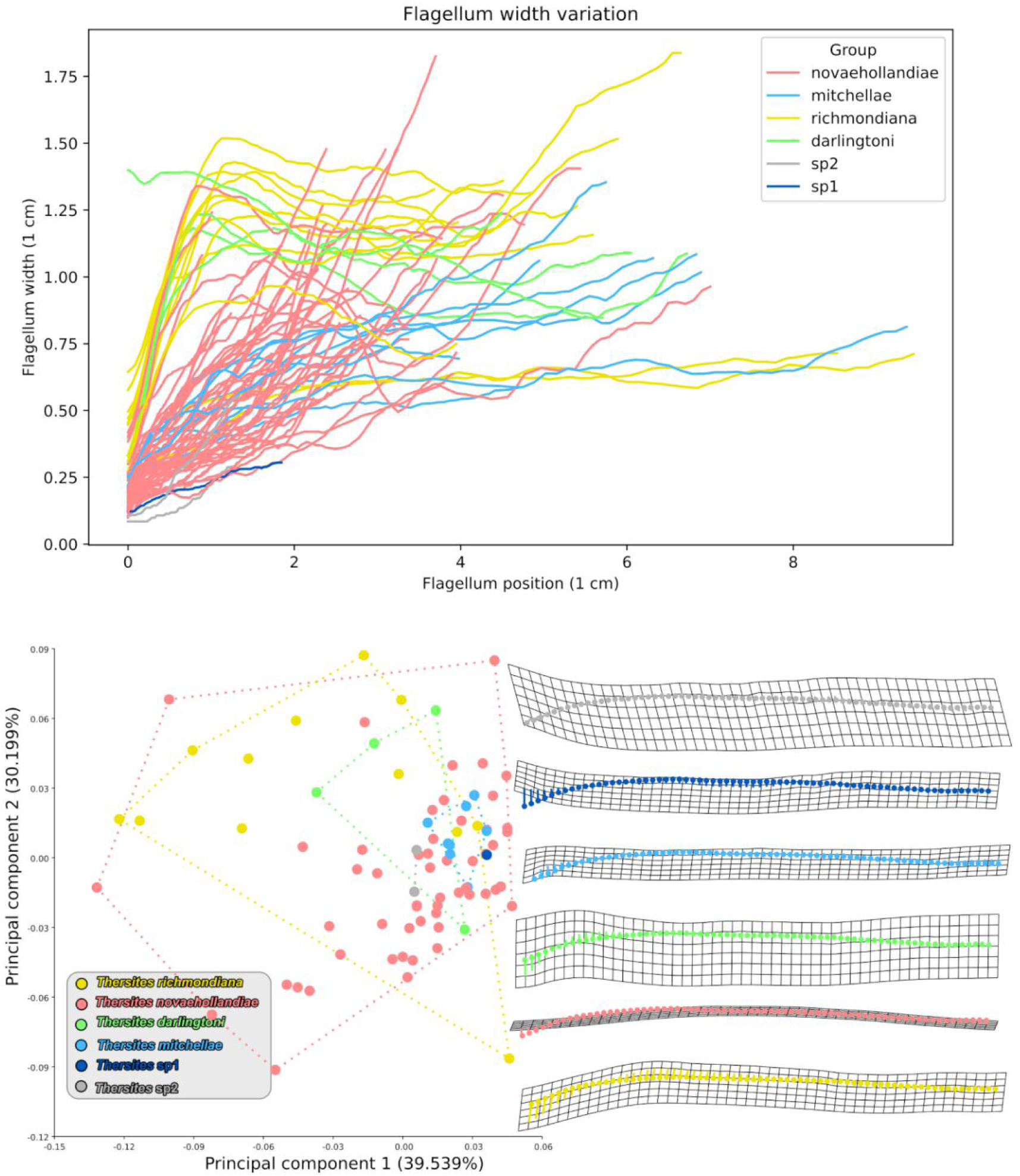
Curves of 70% length flagellum shapes and scatter plots of principal component analysis for 70% length flagellum with thin plate spline grid maps for each species.

**Supplementary Fig. 6.**
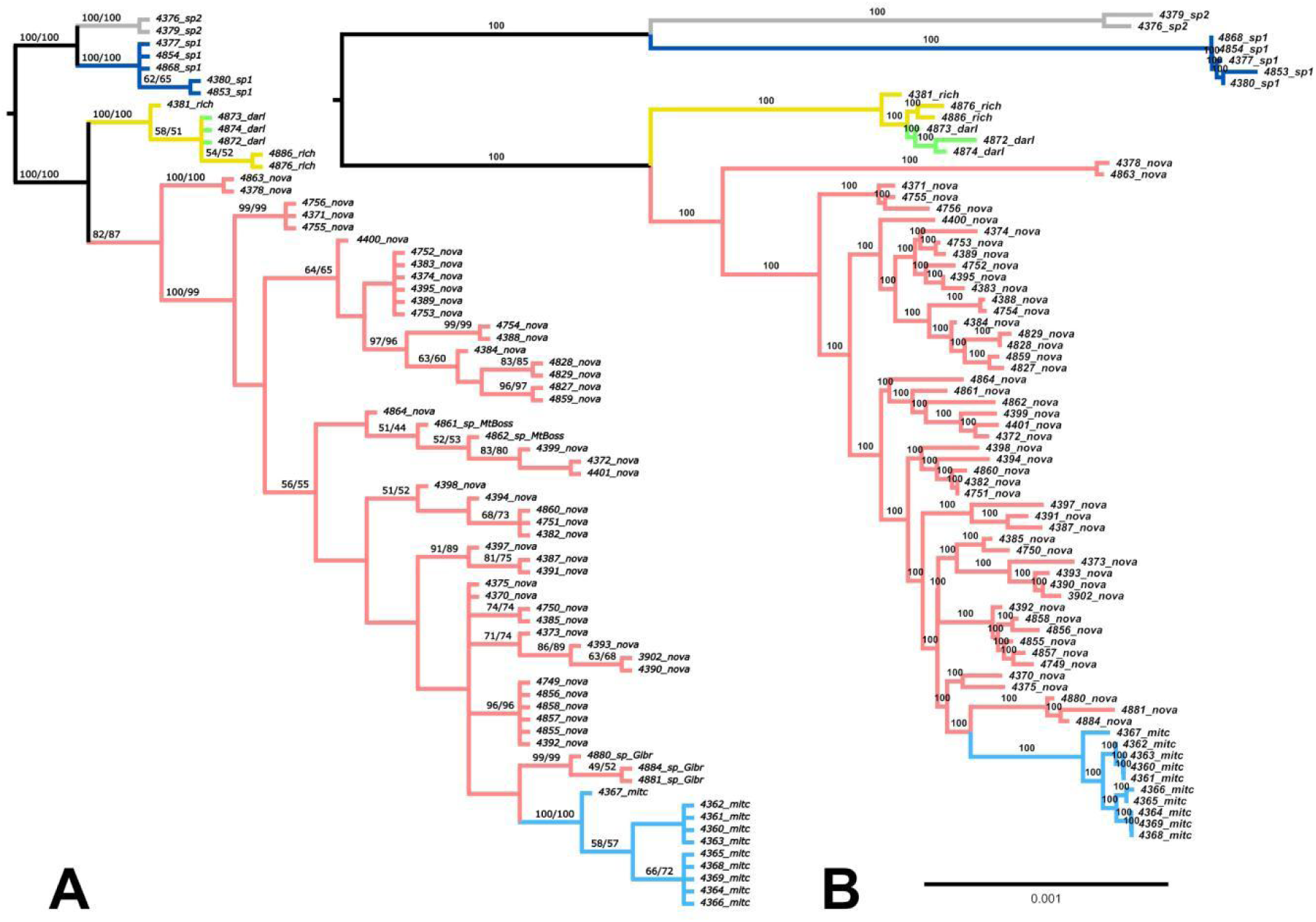
A. Maximum Parsimony tree under equal weighting. The Jackknifing and symmetric resampling were marked on the phylogenetics. B. Maximum likelihood tree with jackknifing.

**Supplementary Fig. 7.**
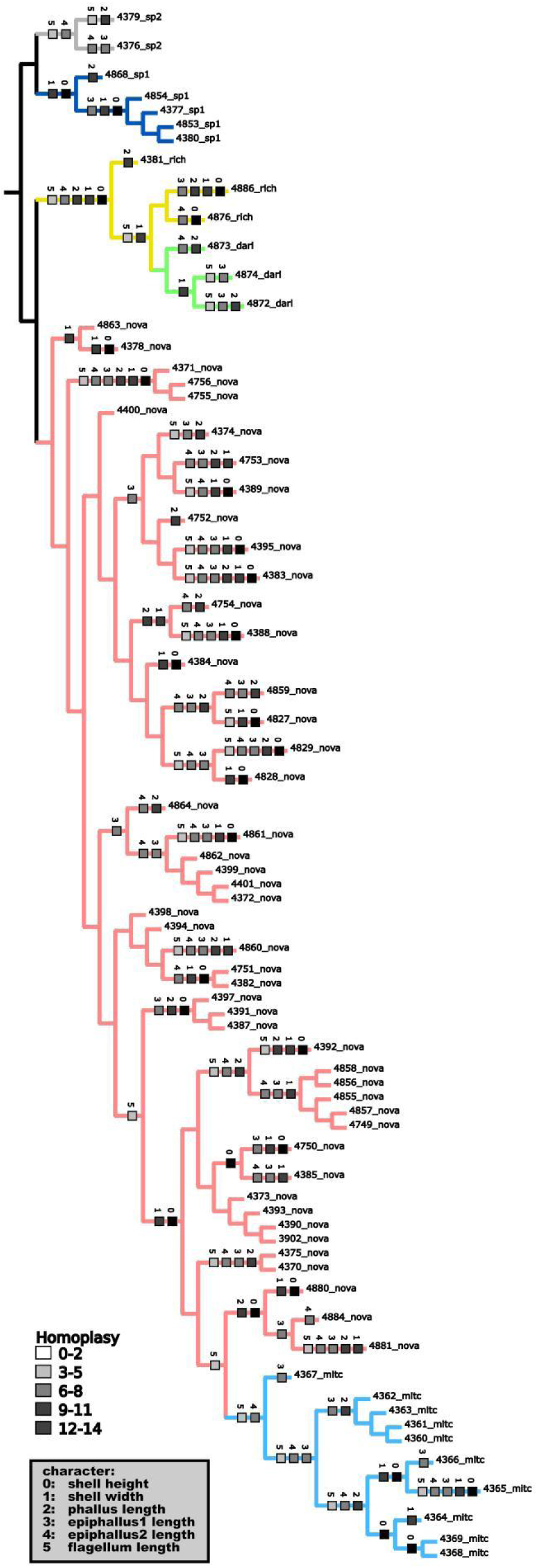
Continuous character measurements optimization. Character homoplasy is indicated on a gradient from white (low) to black (high).

**Supplementary Fig. 8.**
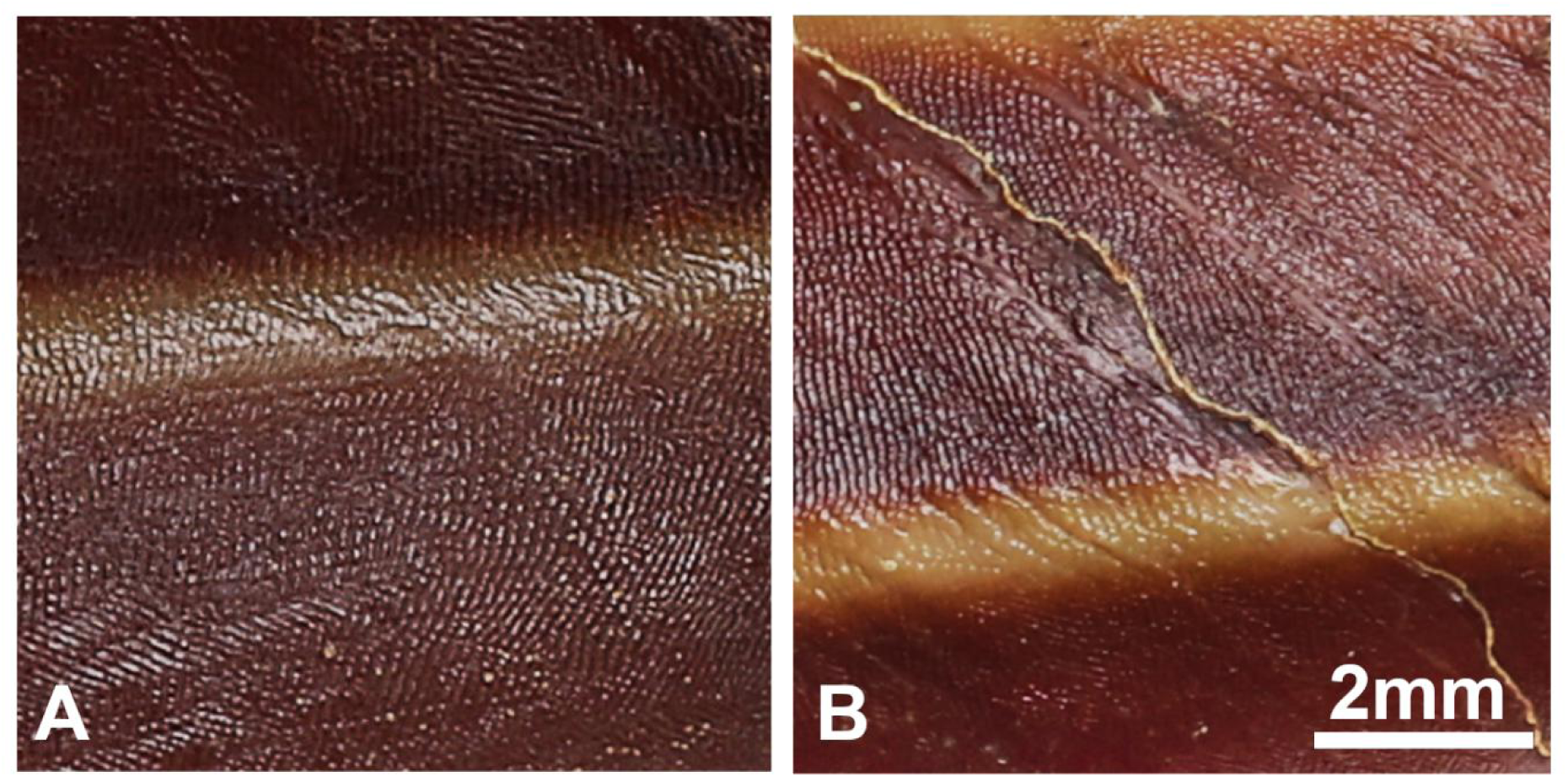
Shell microstructures. A. Fine radial wrinkles; B. Fine radial wrinkles crossed, forming ridgelets. The distinction depends on the proportion of radial wrinkles intersected.

